# Hierarchical regulation of cerebellar neurogenesis by Sin3A-mediated gene repression

**DOI:** 10.1101/2025.10.17.683101

**Authors:** Lei Chen, Ankita Roy, Gregory David, Chin Chiang

## Abstract

Cerebellar granule cells (GCs) are critical for motor and cognitive functions. Lineage tracing studies have identified a hierarchical developmental progression of GC neurogenesis, transitioning from Sox2^+^ stem-like cells to Atoh1^+^ rapidly proliferating granule cell precursors (GCPs), and ultimately to NeuN^+^ mature GCs. However, the molecular mechanisms governing these transitions remain poorly understood. In this study, we identified a transient, slow-cycling progenitor population defined by co-expression of Sox2 and Atoh1. We show that GC maturation depends critically on the repressive function of the Sin3A/Hdac1 complex, which sequentially silences Sox2 and then Atoh1 to ensure orderly progression through developmental stages. Loss of these repressions prolongs progenitor states, compromises survival, and markedly reduces GC output. We also identify NeuroD1 as a co-repressor that collaborates with Sin3A/Hdac1 to inhibit Atoh1 transcription. Our findings highlight the central role of the Sin3A complex in orchestrating distinct stages of cerebellar GC lineage development and may provide insights into Sin3A-related cerebellar disorders and medulloblastoma in human.

**Graphical Abstract:** 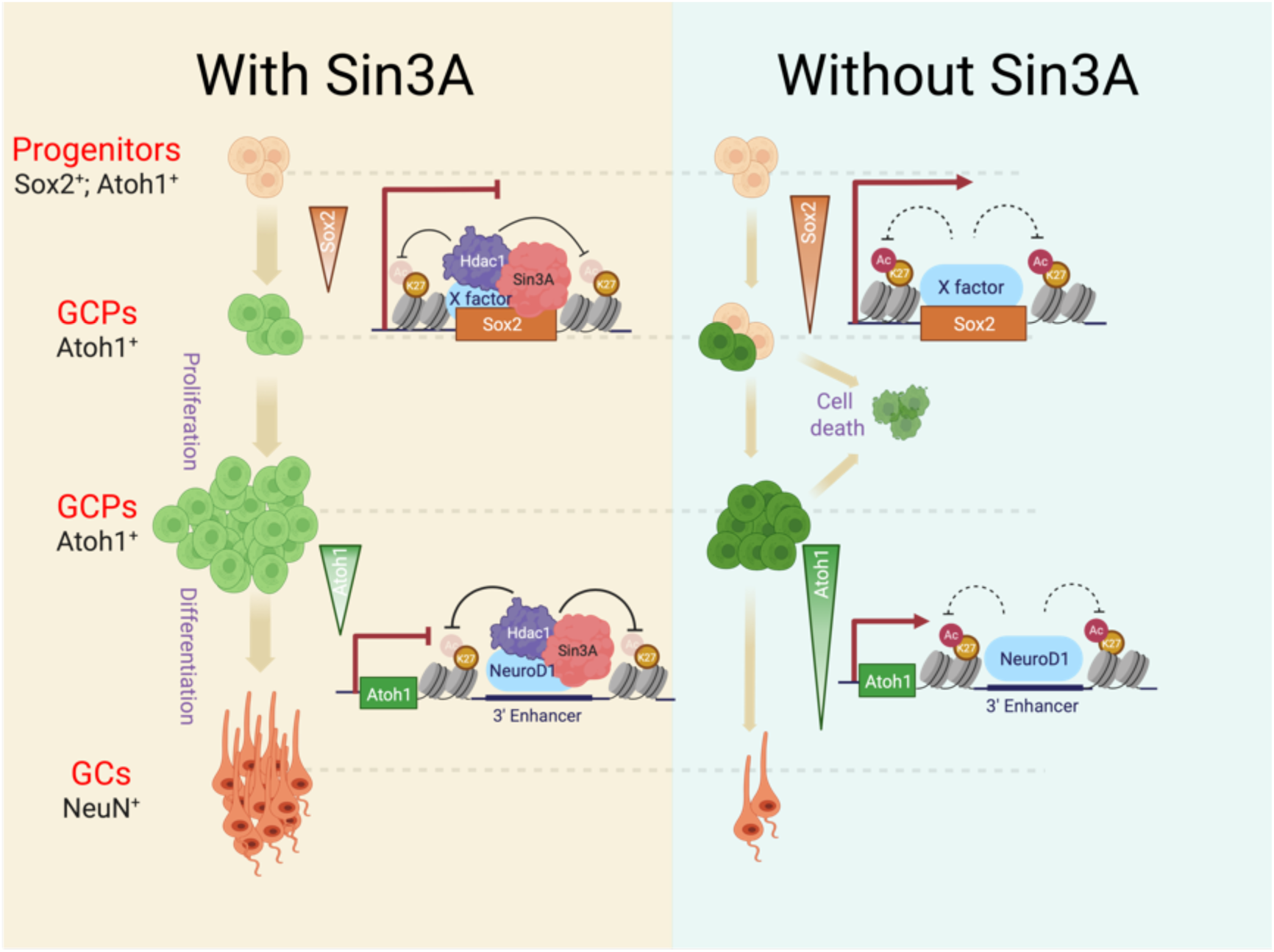

**Highlights:**

- Sin3A is sequentially required for GC lineage progression
- Sin3A promotes the transition of slow-cycling progenitors to GCPs by repressing Sox2 expression
- Sin3A facilitates GCP differentiation by repressing Atoh1 expression
- NeuroD1 recruits the Sin3A/Hdac1 complex to suppress Atoh1 expression

## Background

The neural circuit of the adult brain, including that of the cerebellar system, comprises a defined number of inhibitory and excitatory neurons. The balance between these biochemically and functionally distinct neurons are essential for normal brain function. Cerebellar granule cells (GCs), the most abundant neurons in the brain, are the predominant excitatory neurons^1^ and contribute more than 95% of the cerebellar volume.^2^ They function to integrate and relay sensory,^3–5^ motor,^6^ and cognitive ^7^ information from mossy fibers of different pre-cerebellar origins. GCs originate from a small group of progenitor cells located in the rhombic lip (RL),^8^ a temporary embryonic germinal zone at the dorsal edge of the hindbrain. From progenitors to neurons, they undergo a hierarchical developmental neurogenesis process.^9^ At the top of this hierarchy, Sox2-expressing progenitors maintain self-renewal with relatively slow cell cycles.^9,10^ Around embryonic day 13 (E13), these cells migrate anteriorly to cover the dorsal surface of the cerebellar anlage, forming a transient germinal center known as the external granular layer (EGL).^8,11^ At here, they gradually cease expressing Sox2 and transition into rapidly proliferating granule cell precursors (GCPs).^9,12^ Soon after their last mitosis, immature GCs transiently remain in the inner EGL before initiating radial migration to the inner granular cell layer (IGL), where they complete their terminal differentiation by three weeks postnatally in mice. In humans, the GCs are generated over a period of a year after birth^13,14^ making the process susceptible to a variety of environmental and chemical insults.^15^ Accordingly, cerebellar hypoplasia is one of the most common brain complications in premature infants with poor developmental outcomes^16^ and is the second most significant highest risk factor for autism.^17^

Sox2 is a SOXB1-HMG box transcription factor that serves as a marker widely expressed in neural stem and progenitor cells within the central nervous system.^18,19^ Lineage tracing studies have demonstrated that both excitatory and inhibitory neurons in the cerebellum are derived from Sox2^+^ progenitors.^10^ Sox2 is crucial for maintaining neural stem/progenitor cells in their slow-cycling states.^20–22^ In the developing neocortex, sustained Sox2 expression inhibits the activation of radial glial cells to rapid proliferating progenitors, while reduced Sox2 expression promotes this process.^23,24^ Therefore, timely withdrawal of Sox2 expression is essential for the progression of neuronal lineage. However, the mechanism that regulate the transition of EGL Sox2^+^ cells to GCPs remains unclear.

A master intrinsic regulator of GCP identity is the Atoh1 transcription factor essential for the formation of GC lineage.^25–27^ Transient misexpression of Atoh1 in the ventricular zone of the cerebellum reprograms inhibitory neurons to GC fate.^28^ Overexpression of *Atoh1* inhibits,^27,29^ whereas postnatal depletion of *Atoh1* promotes,^27,30–32^ GCP differentiation. Moreover, Atoh1 cooperates with Gli2, a key downstream mediator of the Shh signaling pathway, as well as promotes Shh responsiveness for the proliferation of GCPs.^30,33^ Therefore, termination of *Atoh1* expression and cell cycle progression is necessary for timely GCP differentiation. However, the mechanisms underlying the coordinated inhibition of *Atoh1* expression to facilitate GCP differentiation remain unclear.

Dynamic changes in gene expression are generally associated with widespread alterations of the epigenetic landscape of the chromatin. Although epigenetic regulators of histone proteins are essential for cerebellar development,^34–36^ whether and how they control different populations of GC lineage fates is unknown. In cells, histone deacetylases(HDACs) forms several distinct and separate multiprotein corepressor complexes with different scaffold proteins that are recruited to various chromatin regions by distinct transcription factors for locus-specific regulation. The core component of one such complex is Sin3A which has shown to associate with HDACs and the transcription factor REST to repress neuronal specific genes in both neuronal and non-neuronal cell lines.^37,38^ Sin3A also regulates stem/progenitor cell maintenance and differentiation.^39–43^ In the developing cortex*, Sin3A* knockdown have shown to reduce cortical progenitor population and alter neuronal differentiation.^42^ Patients with heterozygous mutations in *SIN3A* cause Witteveen-Kolk syndrome, a multisystemic hereditary developmental disorder.^42^ The most prominent phenotype of this condition is a neurodevelopmental disorder characterized by intellectual disability with heterogeneous clinical features such as autism spectrum disorder (ASD) and motor function deficits including cerebellar ataxia,^44–48^ features that are shared with cerebellar dysfunction.

In this study, we reveal the Sin3A repressor complex facilitates the progressive transitions of GC lineage development. Mechanistically, the Sin3A complex epigenetically represses *Sox2* and *Atoh1* expression, permitting timely transitioning from slow cycling progenitors to rapidly dividing GCPs, and finally to differentiated GCs. We further reveal that the Sin3A/Hdac1 complex recruits NeuroD1 and leverages its DNA-binding capability to target the *Atoh1* locus. There, it exerts its deacetylase activity to reduce the levels of H3K27 acetylation at regulatory elements of *Atoh1*, thereby facilitating the timely withdrawal of Atoh1 expression during cellular differentiation.

## Results

### Loss of Sin3A in cerebellar GCPs leads to a diminished GC lineage pool

To understand how Sin3A deficiency leads to cerebellum-associated neurological defects, we generated a loss-of-function mouse model by ablating Sin3A in GCPs. The mouse line harboring the floxed *Sin3A* allele, designated as *Sin3A^Flox^*,^49^ were cross with *Atoh1-Cre* mice that express Cre recombinase in GCPs.^50^ *Sin3A^Flox/Flox^; Atoh1-Cre* (designated here as *Sin3A^CKO^*) mice were born at expected Mendelian ratios. These mice survived into adulthood and exhibited an ataxia-like phenotype, which was characterized by symptoms including tremors and lateral rolling (Supplementary video). The cerebellum of *Sin3A^CKO^* mice at postnatal day 40 (P40) was markedly smaller compared to that of the control group (*Atoh1-Cre* or *Sin3A ^Flox/Flox^* or *Sin3A ^Flox/+^,* designated here as control, Fig. 1A). Morphological analysis revealed a significant reduction in foliation within the anterior region of the cerebellum; lobules Ι to III are represented by a single lobule and lobules IV-VII are hypoplastic (Fig. 1A). Moreover, the characteristic three-layered cortical laminar organization is absent in the most severely affected anterior lobule. In contrast, the posterior lobules VIII to X are relatively normal, which is likely attributed to the incomplete Cre recombinase expression in the posterior lobules as reported previously.^51^ Indeed, immunolabeling of Cre confirmed its expression in anterior EGL (Fig. 1B1 and 1B3), while it was absent in posterior EGL (Fig. 1B2 and 1B4), corroborated by the more efficient Sin3A deletion in the anterior GCPs (Fig. S1A, panels A3 and A4). This incomplete deletion serves as an internal control to examine how Sin3A regulates GCP proliferation and differentiation by comparing the cells in the anterior (I to VII) and posterior (VIII to X) regions.

**Figure 1.**
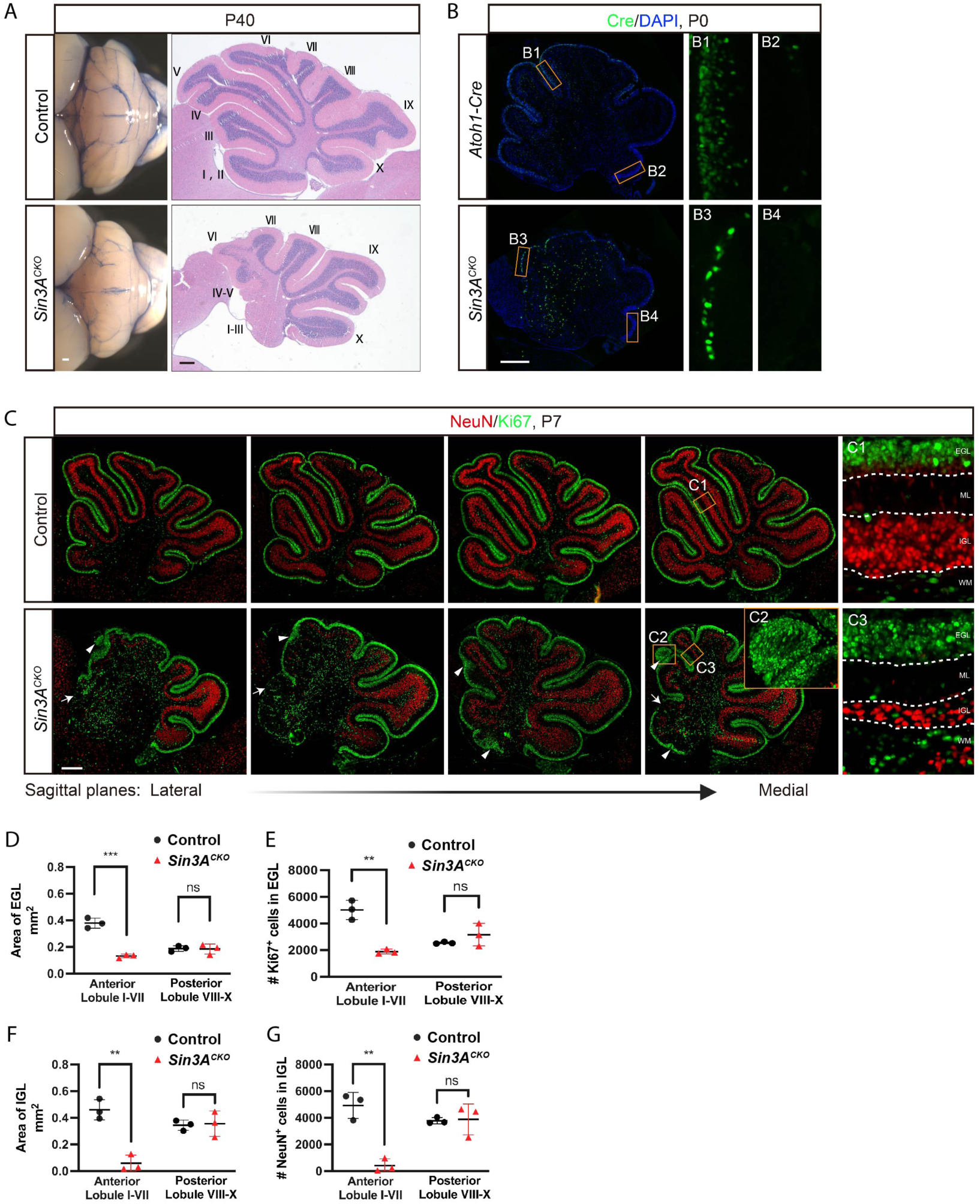
Loss of Sin3A in cerebellar GCPs leads to a diminished GC lineage pool. (A) External views (left) and H&E stained midsagittal sections (right) from control and *Sin3A^CKO^* mice at postnatal day 40 (P40), showing a reduction in cerebellar size and foliation. Scale bars indicate 250 μm. (B) Immunofluorescence staining for Cre recombinase at P0 in the cerebellum of control and *Sin3A^CKO^* mice. Nuclei were counterstained with DAPI. Scale bar indicates 250 μm. (C) Immunofluorescence staining for Ki67 and NeuN at P7 in the cerebellum of control and *Sin3A^CKO^* mice. A series of sections from the lateral to medial cerebellum revealing significant variability in *Sin3A^CKO^* across different sagittal planes, attributable to disorganization within the tissue architecture. The dashed lines delineate the boundaries of different layers within the cerebellar cortex. EGL: external granular layer, ML: molecular layer, IGL: internal granular layer, PWM: prospective white matter. Arrows indicate acellular gaps in EGL. Arrowheads indicate ectopic clusters of GCPs. Scale bar indicates 250 μm. (D and E) Quantification of the cross-sectional area of EGL (D) and the number of Ki67^+^ cells (E) in EGL. Shown are mean ± SEM. Two-tailed t test: ** p ≤ 0.01, *** p ≤ 0.001, n.s. p>0.05. n= 3 mice per group. (F and G) Quantification of the cross-sectional area of IGL (F) and the number of NeuN^+^ cells (G) in IGL. Shown are mean ± SEM. Two-tailed t test: ** p ≤ 0.01, *** p ≤ 0.001, n.s. p>0.05. n= 3 mice per group.

We next focused on the critical period of GCP development, which occurs within the first three weeks postnatally. At the peak of GCP proliferation in P7, we investigated the developmental progression of the GC lineage by employing immunolabeling to detect the proliferation marker Ki67 and the terminal differentiation marker NeuN. In the control mice, a characteristic four-layered structure representing EGL, molecular layer (ML), internal granule layer (IGL) and white matter (WM) was observed (Fig. 1C, panel C1). Within this structure, the GC lineage was densely distributed across two distinct layers: the EGL, which is composed of Ki67^+^ GCPs and weak NeuN^+^ immature GCs, and the IGL, characterized by strong NeuN expression indicative of mature GCs. In contrast, the anterior region of *Sin3A^CKO^*cerebella largely failed to form a well-organized cellular layering and exhibited high variability across different sagittal sections (Fig. 1C). Notably, a continuous EGL was not observed and interrupted by acellular gaps (arrows in Fig. 1C). Accordingly, there was a significant decrease in the total area of EGL (Fig. 1D) and the number of GCPs (Fig. 1E). Despite this reduction, we observed reginal thickening of the EGL, with occasional ectopic clusters of GCPs (arrowheads in Fig. 1C, panel C2). NeuN labeling revealed a pronounced decrease in mature GCs in the IGL (Fig. 1F and 1G). Interestingly, the reduction of GCs is observed even in the regions where the thickness of EGL is comparable to the control (Fig. 1C, panel C1 and C3), suggesting differentiation of GCPs is compromised.

### GCPs fail to differentiate and undergo apoptosis in the absence of Sin3A

There are three mutually non-exclusive potential causes for the observed reduction of GC lineage: 1) impaired proliferation of GCPs, 2) disrupted differentiation of GCPs, and 3) increased cell death. To address the first possibility, we assessed the relative abundance of proliferating GCPs at two developmental stages. At P0 when distinct lobules begin to emerge, the percentage of Ki67^+^ GCPs in the EGL of *Sin3A^CKO^* cerebella remains similar to that of controls, albeit the total number of GCPs is much reduced (Fig. 2A, 2B and Fig. S2A, S2B). By P7, the proportion of Ki67^+^ GCPs in the EGL is significantly higher in *Sin3A^CKO^* mice (Fig. 2A and 2B). To further analyze the proliferation status of GCPs, we assessed the S-phase fraction, which serves as an indicator of proliferative activity.^52,53^ One hour before euthanasia, mice received an injection of 5-ethynyl-2’-deoxyuridine (EdU), a thymidine analogue that labels cells in the S phase of the cell cycle. Consistent with above proliferation data, the fraction of GCPs in S phase (EdU^+^; Ki67^+^ cells/ Ki67^+^ cells) was significantly higher in the P7 stage *Sin3A^CKO^*cerebellum compared to control (Fig. 2C). We also assessed the expression of Cyclin D1 (Ccnd1), a cell cycle regulator expressed in the proliferating cells, and found that the percentage of Ccnd1+ cells is also increased, supporting the observation that *Sin3A^CKO^* GCPs exhibit increased proliferative activity rather than entering a state of cell cycle arrest (Fig. S2C). Therefore, the observed reduction in the GC population cannot be attributed to decreased GCP proliferation.

**Figure 2.**
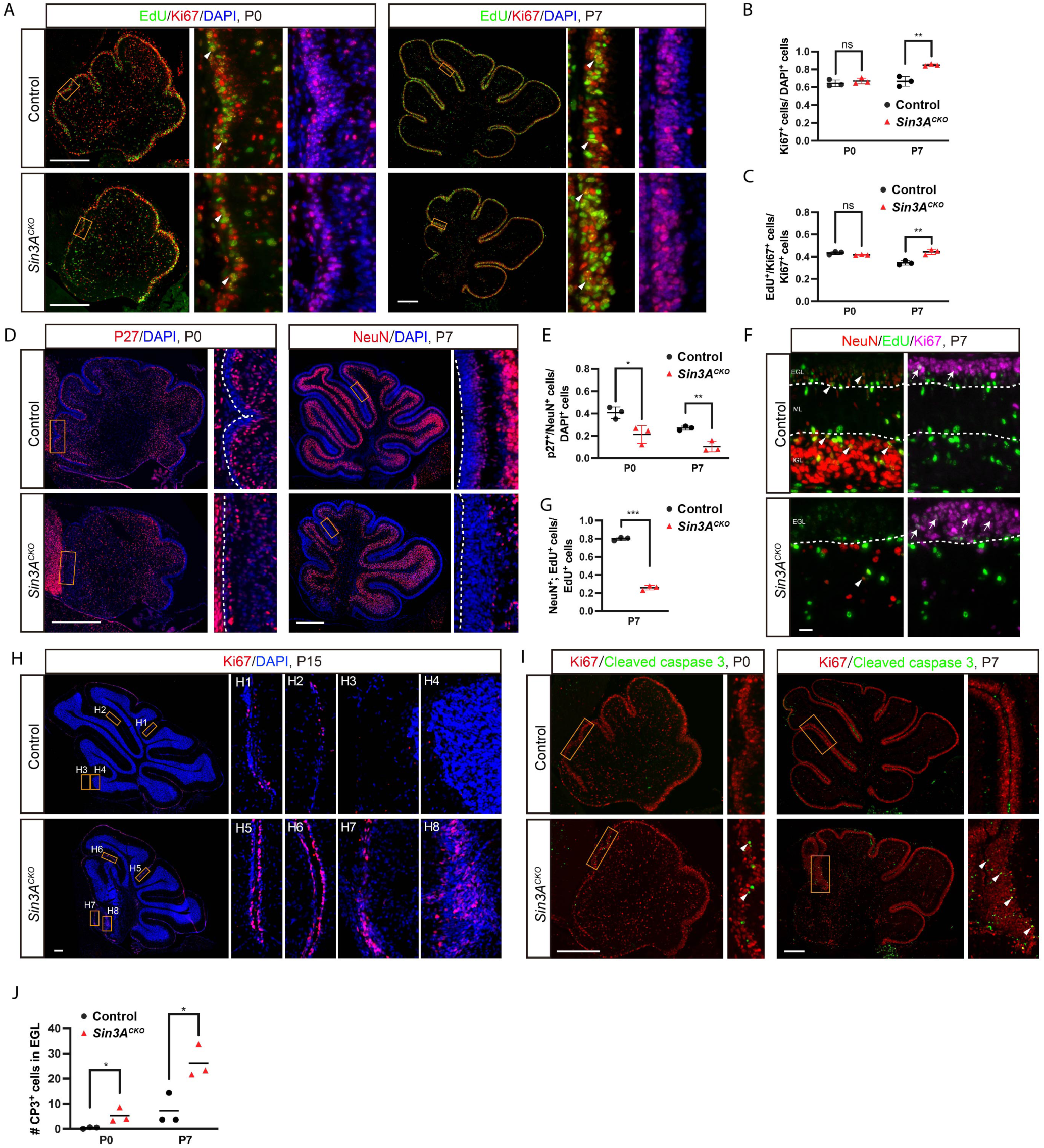
GCPs fail to differentiate and undergo apoptosis in the absence of Sin3A. (A) Ki67 and EdU staining in cerebellum of control and *Sin3A^CKO^* mice at P0 and P7. Nuclei were counterstained with DAPI. Mice were administrated with EdU 30 minutes before sacrifice. Arrowheads indicate cells positive for both Ki67 and EdU. Scale bars indicate 250 μm. (B) Quantification of the percentage of Ki67^+^ cells relative to the total number of DAPI^+^ cells in EGL at P0 and P7. Shown are mean ± SEM. Two-tailed t test: ** p ≤ 0.01, n.s. p>0.05. n= 3 mice per group. (C) Quantification of the percentage of EdU^+^/Ki67^+^ double-positive cells relative to the total number of Ki67^+^ cells in EGL at P0 and P7. Shown are mean ± SEM. Two-tailed t test: ** p ≤ 0.01, n.s. p>0.05. n= 3 mice per group. (D) Immunofluorescence staining for p27 at P0 and NeuN at P7 in the cerebellum of control and *Sin3A^CKO^*mice. Nuclei were counterstained with DAPI. The dashed lines delineate the outer margin of the EGL. Scale bars indicate 250 μm. (E) Quantitative analysis showing the percentage of p27^+^ or NeuN^+^ cells relative to the total cell population in the EGL. Shown are mean ± SEM. Two-tailed t test: * p ≤ 0.05, ** p ≤ 0.01. n= 3 mice per group. (F) Ki67, NeuN and EdU staining in the cerebellum of control and *Sin3A^CKO^* mice at P7. Nuclei were counterstained with DAPI. Mice were administrated EdU 48 hour before sacrifice. Arrowheads indicate cells positive for both NeuN and EdU, and arrows indicate cells positive for both Ki67 and EdU. The dashed lines delineate the boundaries of different layers within the cerebellar cortex. Scale bars indicate 250 μm. (G) Quantification of the percentage of EdU^+^/NeuN^+^ double-positive cells relative to the total number of EdU^+^ cells. Shown are mean ± SEM. Two-tailed t test: *** p ≤ 0.001. n= 3 mice per group. (H) Immunofluorescence analysis for Ki67 at P15 in the cerebellum of control and *Sin3A^CKO^* mice. Nuclei were counterstained with DAPI. Scale bars indicate 250 μm. (I) Immunofluorescence staining for cleaved caspase-3 (CP3) and Ki67 at P0 and P7 in the cerebellum of control and *Sin3A^CKO^* mice. Arrowheads indicate CP3^+^ cells. Scale bars indicate 250 μm. (J) Quantification of the CP3^+^ cells in the EGL. Shown are mean ± SEM. Two-tailed t test: * p ≤ 0.05. n= 3 mice per group.

To assess whether *Sin3A* depletion affects GCP differentiation, we analyzed the proportion of differentiated cells relative to the total cell population in the EGL. Through immunolabeling with the differentiated cell marker Cdkn1b (p27), we found that at the onset of postnatal development (P0), the innermost 2-3 cell layers of the EGL in control mice had initiated differentiation and were progressively migrating out of the EGL. Notably, in *Sin3A^CKO^*, only a sparse number of cells exhibited p27^+^ (Fig. 2D), with a significantly lower proportion of differentiated cells compared to control (Fig. 2E). Similar differentiation defects were observed at P7 when NeuN was used as a more specific marker for GCs (Fig. 2E). We then traced the differentiation potential of GCPs by labeling them with EdU at P5, and determined their NueN+ descendants 48 hours later. The results showed that in the control cerebellum, over 80% of EdU^+^ cells differentiated and migrated inward to the inner EGL or the IGL within this timeframe (indicated by arrowheads in Fig. 2F), whereas only about 25% did so in the *Sin3A^CKO^* (Fig. 2F and 2G). GCPs that had not differentiated remained in the Ki67^+^ zone of the EGL, exhibiting diminished EdU staining intensity due to clonal dilutions as a result of ongoing cell division (indicated by arrows in Fig. 2F). In regions with comparable EGL thickness, dual labeling of Ki67 and NeuN clearly showed an enlargement of the proliferative cell zone alongside a diminished differentiated cell layer in *Sin3A^CKO^* (Fig. S2D). Moreover, a large number of GCPs persist in *Sin3A^CKO^* at P15 (Fig. 2H, panel H5-H8), even as the vast majority of GCPs have already differentiated and migrated away from the EGL in the control (Fig. Fig. 2H, panel H1-H4). These results indicate that in the absence of Sin3A function, differentiation of GCPs is markedly inhibited, resulting in their retention in a proliferative state. However, the total GCP pool remains small, suggesting some of them are actively eliminated.

To determine if cell death contribute to the reduction of GCPs in *Sin3A^CKO^* mice, we employed cleaved caspase-3 (CP3) immunostaining, and found that the number of apoptotic GCPs in *Sin3A^CKO^* mice was significantly higher than that in control mice both at P0 and P7 (arrowheads Fig. 2I and 2J). Moreover, immunostaining with γ-H2AX, a marker for DNA damage, revealed double-stranded DNA breaks (DSBs) in a subset of GCPs from *Sin3A^CKO^* mice (Fig. S2E). The cell death is specific to GCPs as we did not detect significant apoptosis in mature GCs (Fig. S2F). In addition, we determine the fate of GCPs using *R26R-tdTomato* (*Ai9*) reporter gene.^54^ The activity of Atoh1-Cre within GCPs can activate constitutive expression of tdTomato, thereby serving as a developmental tracer for the GC-lineage cells. Results indicated that in the anterior parts of the *Sin3A^CKO^* cerebellum, only limited number of tdTomato^+^ cells differentiate into GCs. These cells migrated inward and formed a dysplastic IGL (Fig. S2G). No abnormal increase of tdTomato^+^ cells was observed in surrounding tissues, ruling out aberrant migration of GCPs. Collectively, our findings indicate that Sin3A is essential for GCPs differentiation and survival.

### Sin3A targetome reveals candidate genes associated with phenotypic defects

To analyze the molecular mechanisms by which Sin3A regulates GC lineage development, we conducted single-cell transcriptomic analysis (scRNA-seq) on two sets of control and *Sin3A^CKO^* cerebellar samples at P6 using the 10x Genomics Flex Gene Expression assay (Fig. 3A). After filtering, transcriptome data from 31,661 cells were recovered, with 16,223 cells originating from control and 15,438 cells from *Sin3A^CKO^* cerebellum. Through cell clustering and marker gene annotation, a total of 17 distinct cell types were identified, encompassing the majority of cell types found in the cerebellum (Fig. 3B). Notably, based on the expression of *Atoh1*, we identified two adjacent cell clusters: GCP-1 and GCP-2 (Fig. S3A). Cell cycle analysis revealed significant differences in phase distribution between these two clusters (Fig. S3A), suggesting that their divergence primarily result from differential expression of cell cycle-related genes. This indicates that future analyses of the GCP population should include regression correction for cell cycle effects. By splitting cells from different genotypes in the UMAP visualization (Fig. S3B), we did not observe any gain or loss of specific cell types in the cerebellum of *Sin3A^CKO^*. However, there were noticeable differences in cell abundance within certain clusters (Fig. S3B). This indicates that the absence of *Sin3A* affects transcriptional characteristics at the sub-cluster level. Therefore, we conducted a quantitative analysis of the differences in abundance using k-nearest neighbor (KNN) graphs.^55^ All cells were assigned into 2,562 partially overlapping neighborhoods based on KNN (Fig. 3C). Statistical analysis revealed that 208 of these neighborhoods showed significant differences (Fig. 3C and 3D). Consistent with differentiation defects noted above, there was a significant reduction in postmitotic GCPs and mature GCs in the *Sin3A^CKO^*cerebellum, while proliferative state GCPs were significantly enriched (Fig. 3C and 3D).

**Figure 3.**
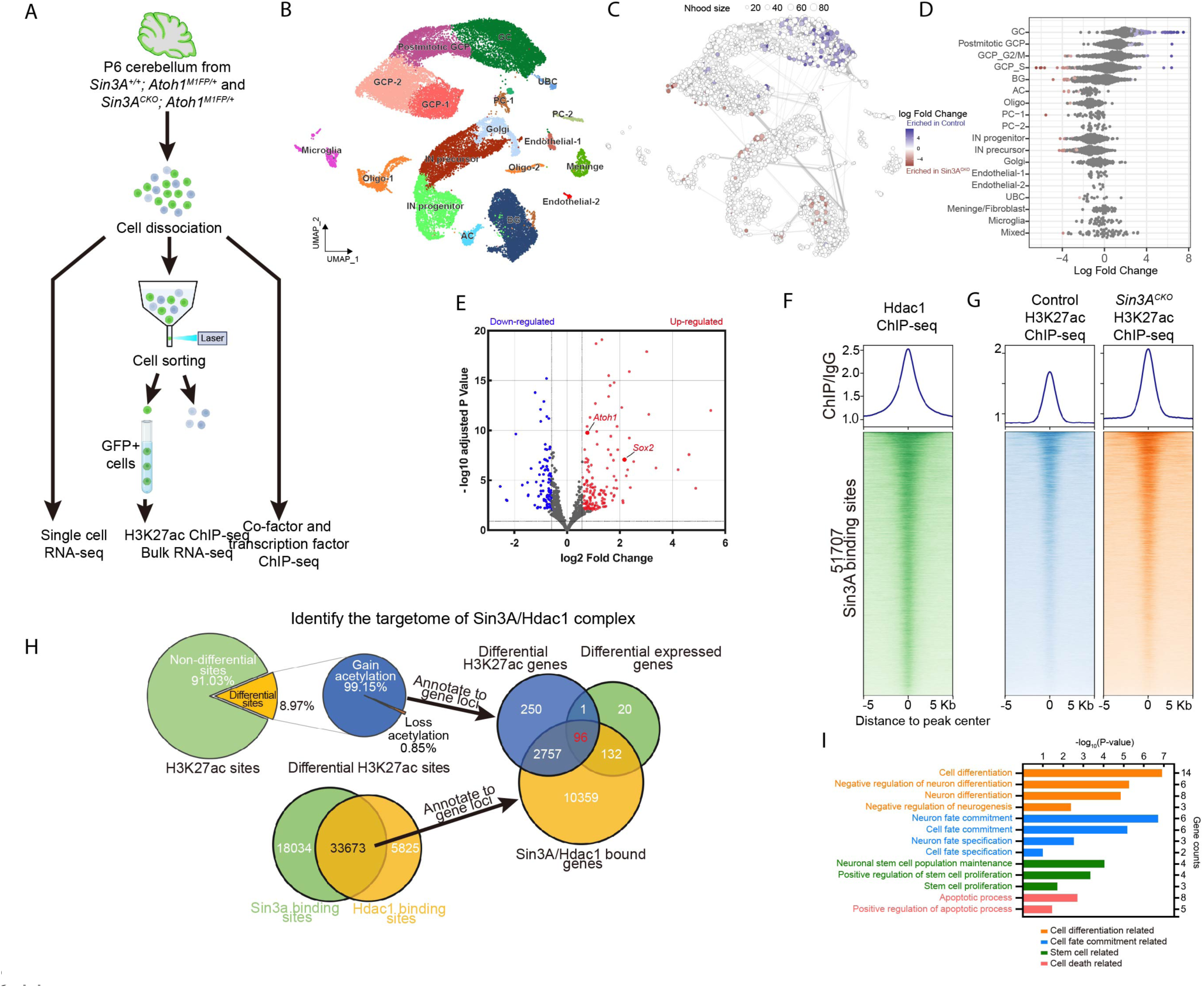
Sin3A targetome reveals candidate genes associated with phenotypic defects. (A) Schematic representation of the strategy employed to isolate cells followed by high-throughput sequencing analysis. (B) Uniform Manifold Approximation and Projection (UMAP) visualization of cell populations from the P6 cerebellum of two *Sin3A^CKO^* and two control mice. UBC: unipolar brush cells, PC: purkinje cells, Oligo: oligodendrocytes, IN: GABAergic interneurons, Golgi: golgi cells, AC: astrocytes, BG: bergmann glial cells, (C) UMAP representation of the results from Milo differential abundance (DA) testing between *Sin3A^CKO^* and control cerebellum with nodes showing cell neighborhoods. Nodes with significant differential abundance are highlighted using a gradient scale of red or blue. (D) Beeswarm plot showing the log-fold change between *Sin3A^CKO^* and control cerebellum across groups of nearest-neighbor cells from various cell type clusters. Significant changes are highlighted using a gradient scale of red or blue. (E) Volcano plot illustrating the differential gene expression results between the GCP clusters of control and *Sin3A^CKO^* cerebellum. (F) Heatmap and average intensity plots of the Hdac1 ChIP-seq signals across 10 kb from the center of Sin3A peaks throughout the genome the genome from dissociated cerebellar cells of control mice. (G) Heatmap and average intensity plots of the H3K27ac ChIP-seq signals across 10 kb from the center of Sin3A peaks throughout the genome from isolated GCPs of control and *Sin3A^CKO^* mice. (H) Pie chart and Venn diagram illustrating the process of screening and identifying target genes of Sin3A. (I) Bar chart showing the results of the functional enrichment analysis for 96 candidate target genes.

Since the deletion of *Sin3A* activated by Atoh1-Cre directly affects GCPs, we subsequently subsetted GCPs and conducted differential expression analysis between different genotypes. We identified 249 differentially expressed genes between *Sin3A^CKO^* and control GCPs, with 150 genes upregulated and 99 downregulated (Fig. 3E, Supplementary Table 1). Additionally, we validated these differential expressions using bulk RNA-seq, 42.5% (106/249) of them were also found to be differentially expressed in the bulk RNA-seq data of P6 GCPs. We then asked which of these differential expressed genes are bound by Sin3A using genome wide chromatin immunoprecipitation followed by deep sequencing (ChIP-seq) analysis. The results identified 51,707 peaks associated with 15,008 genes. We then cross-referenced to 249 genes selected from scRNA-seq and identified 233 direct target genes of Sin3A repression. As a corepressor, Sin3A forms complexes with HDACs in cells, thereby regulating the activity of target gene cis-regulatory elements (CREs) through deacetylation of histone lysine residues including H3K27ac and consequently modulating gene transcription.^56,57^ We therefore performed Hdac1 ChIP-seq analysis and found that the genome-wide binding profiles of Hdac1 and Sin3A are highly comparable (Fig. 3F), with 65.1% of Sin3A peaks overlapping with that of Hdac1 (Fig. 3H). This is expected given that Hdac1 is also known to form Sin3A-independent complexes.^58^ Notably, 97.8% of the Sin3A target genes (228/233) have overlapping binding sites between Sin3A and Hdac1. We next assessed how many of these 228 Sin3A/Hdac1 target genes displayed enhanced acetylation of H3K27 in *Sin3A^CKO^*GCPs. If the Sin3A/Hdac1 complex canonically suppresses gene expression in GCPs through deactivating CREs, we would expect a genome-wide hyperacetylation of H3K27 in Sin3A null GCPs. To circumvent the potential confounding effects of other cell types, we employed *Atoh1^M1GFP^*, a knock-in mouse line with a fused EGFP to the C-terminus of Atoh1 protein,^59^ for the fluorescence-activated cell sorting purification of GCPs (Fig. 3A). After H3K27ac ChIP-seq with purified GCPs, we identified a 47% increase in H3K27ac enrichment at global Sin3A binding sites within *Sin3A^CKO^*GCPs when compared to the control (Fig. 3G). In the genome of GCPs, among 48,445 H3K27ac-enriched loci, approximately 8.97% exhibited differential acetylation following *Sin3A* depletion. Notably, the vast majority (99.15%) of these differentially acetylated sites showed an increase in H3K27ac levels (Fig. 3H). The enriched peaks were mapped to 3104 genes and of which 96 genes overlapped with Sin3A and Hdac1 target genes (Fig. 3H, Supplementary table 2). The function annotations of these genes are enriched in gene ontology terms related to cell differentiation and apoptosis (Fig. 3I), which is consistent with phenotypic alterations observed in *Sin3A^CKO^* GCPs. Notably, enrichments were also observed in genes associated with stem cell function and cell fate commitment (Fig. 3I), including Sox2 and Atoh1.

### Sox2^+^; Atoh1^+^ cells in the EGL represent slow-cycling and intermediate GCPs

Previous studies have reported that Sox2^+^ cells in the EGL originate from Sox2^+^ embryonic cerebellar progenitors (ECPs) in the RL^10^ and they contribute significantly to the mature granule cells.^9,12^ To better understand Sox2^+^ cells in relation to Atoh1^+^ GCPs in the GC lineage, we analyzed Sox2 and Atoh1 expression at E13.5, a developmental stage during which the GC lineage is undergoing specification and the EGL is initially forming. In the RL of cerebellar primordium, which is enriched with ECPs, nearly all cells express Sox2 (Fig. S4A). Scattered among these are some Sox2^+^; Atoh1^+^ cells, representing a population recently committed to the GC lineage while still maintaining high levels of Sox2 expression (arrows in Fig. S4A). As GC lineage cells migrate, those that have reached the dorsal surface (uRL) downregulate Sox2 to a low level of expression (arrowheads in Fig. S4A). This Sox2^+^; Atoh1^+^ co-expression pattern extends throughout the EGL along GCPs’ tangential migration path (Fig. 4A). Quantitative analysis indicates that over 90% of Atoh1^+^ cells in the EGL co-express Sox2 but lack NeuroD1 expression (Fig. 4E), suggesting they represent a relatively primitive GCPs (arrowheads in Fig. 4A). By E16.5, EGL cells progressively cease expressing Sox2, reducing their proportion among GCPs to 13.8% (Fig. 4E). The remaining Sox2^+^; Atoh1^+^ cells occupy one to two outer layers within the EGL (Fig. 4B2). Beneath them, in the inner layer, are Atoh1^+^; NeuroD1^+^ cells that have already ceased expressing Sox2 (Fig. 4B2). Along the anteroposterior axis of the cerebellum, Sox2 expression gradually decreases and nearly disappears entirely in the most anterior part of the EGL, leaving only Atoh1^+^ GCPs (Fig. 4B1). Given that anterior GCPs represent a population that has initiated migration and settlement earlier,^60^ this wave-like withdrawal of Sox2 from anterior to posterior suggests that, after committing to the GC lineage from ECPs, GCPs transiently maintain Sox2 expression until they complete tangential migration. Accordingly, the proportion of Sox2^+^; Atoh1^+^ cells drastically reduced to 2.5% at P0, and by P7, they are rarely detectable (Fig. 4C1, 4C2, 4D and 4E). Thus, it appears that upon initiating radial migration, these cells gradually cease expressing Sox2 and transition into rapidly proliferating Atoh1^+^ GCPs (Fig. 4F). To support this notion, we analyzed publicly available scRNA-seq data from wild-type mice at E12–E16.^61^ By examining the expression of marker genes across different developmental stages of the GC lineage, such as *Sox2*, *Atoh1*, and *Cntn2*, we were able to subset and finely cluster GC lineage cells (Fig. 4H). Defining *Sox2^+^* ECPs as the starting point of development allowed us to construct a developmental trajectory from progenitors to post-mitotic cells (Fig. 4G). During the transition phase from ECPs to GCPs, we identified a population co-expressing both *Sox2* and *Atoh1*, which represent Sox2^+^; Atoh1^+^ cells described above (Fig. 4H). Consistently, this group constitutes 13.9% of the GCP population at stage E16 (Fig. S4C). Compared to other GCP sub-clusters, Sox2^+^; Atoh1^+^ cells exhibit higher expression of genes associated with neuronal stem cell maintenance and proliferation, such as *Hes1*, *Sox2*, *Fgfr1*, and *Id2*. Conversely, genes related to neurogenesis and neuron differentiation—such as *Nrn1*, *Tubb3*, *NeuroD1*, and *Gap43*—are expressed at lower levels (Fig. S4F and S4G). Furthermore, they have a higher potency score (Fig. S4D), suggesting that these cells are in a relatively primitive state. Indeed, A 30-minute EdU pulsing assay indicated that the proliferative activity of Sox2^+^; Atoh1^+^ cells was significantly lower than that of other GCPs (Fig. 5D and 5E). Thus, Sox2^+^; Atoh1^+^ cells represent a transitory slow-cycling intermediate progenitors of GCPs.

**Figure 4.**
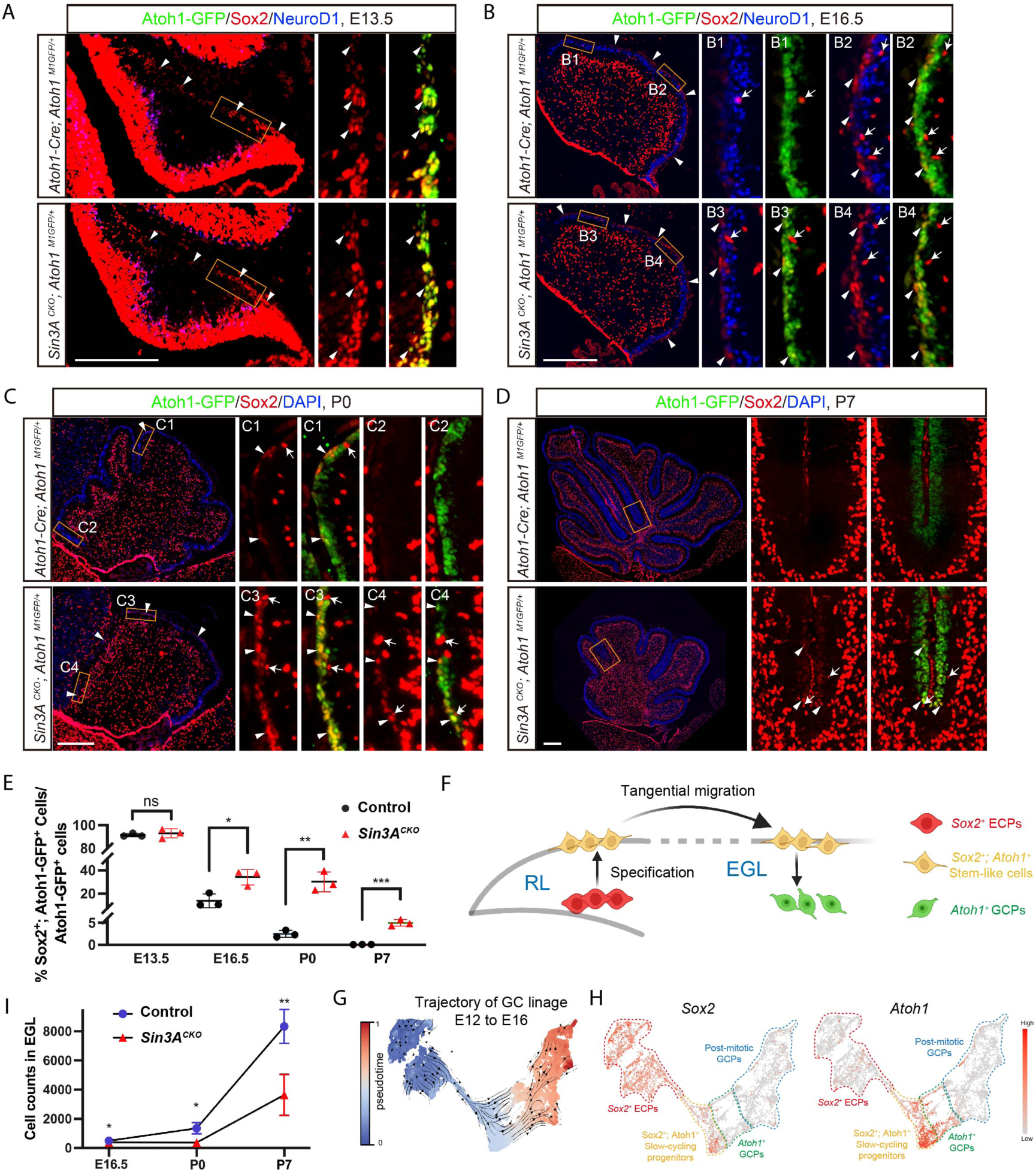
Sox2^+^; Atoh1^+^ cells in the EGL represent slow-cycling and intermediate GCPs. (A) Immunofluorescence staining for Sox2, Atoh1-GFP, and NeuroD1 at E13.5 in the cerebellum of control and *Sin3A^CKO^* mice. Arrowheads indicate cells positive for Sox2 and Atoh1 in the EGL. Scale bar indicates 250 μm. (B) Immunofluorescence staining for Sox2, Atoh1-GFP, and NeuroD1 at E16.5 in the cerebellum of control and *Sin3A^CKO^* mice. Arrowheads indicate cells positive for Sox2 and Atoh1, and arrows indicate cells positive for only Sox2. Scale bar indicates 250 μm. (C) Immunofluorescence staining for Sox2 and Atoh1-GFP at P0 in the cerebellum of control and *Sin3A^CKO^*mice. Nuclei were counterstained with DAPI. Arrowheads indicate cells positive for Sox2 and Atoh1, and arrows indicate cells positive for only Sox2. Scale bar indicates 250 μm. (D) Immunofluorescence staining for Sox2 and Atoh1-GFP at P7 in the cerebellum of control and *Sin3A^CKO^*mice. Nuclei were counterstained with DAPI. Arrowheads indicate cells positive for Sox2 and Atoh1, and arrows indicate cells positive for only Sox2. Scale bar indicates 250 μm. (E) Quantification of the Sox2^+^; Atoh1-GFP^+^ cells relative to the Atoh1-GFP^+^ population in the medial and anterior EGL. Shown are mean ± SEM. Two-tailed t test: * p≤ 0.05, ** p ≤ 0.01, *** p ≤ 0.001, n.s. p>0.05. n= 3 mice per group. (F) Diagram illustrating the developmental model transitioning from multipotent ECPs to rapidly proliferating GCPs. (G) UMAP displaying the developmental trajectory of GC lineages from E12 to E16 wildtype mice, based on pseudotime analysis. (H) UMAP displaying the *Sox2* and *Atoh1* expression in the GC lineage cell of E12 to E16 wildtype mice. (I) Quantification of cells counts in the medial and anterior EGL of E16.5, P0 and P7 cerebellum. Shown are mean ± SEM. Two-tailed t test: * p≤ 0.05, ** p ≤ 0.01. n= 3 or 4 mice per group.

**Figure 5.**
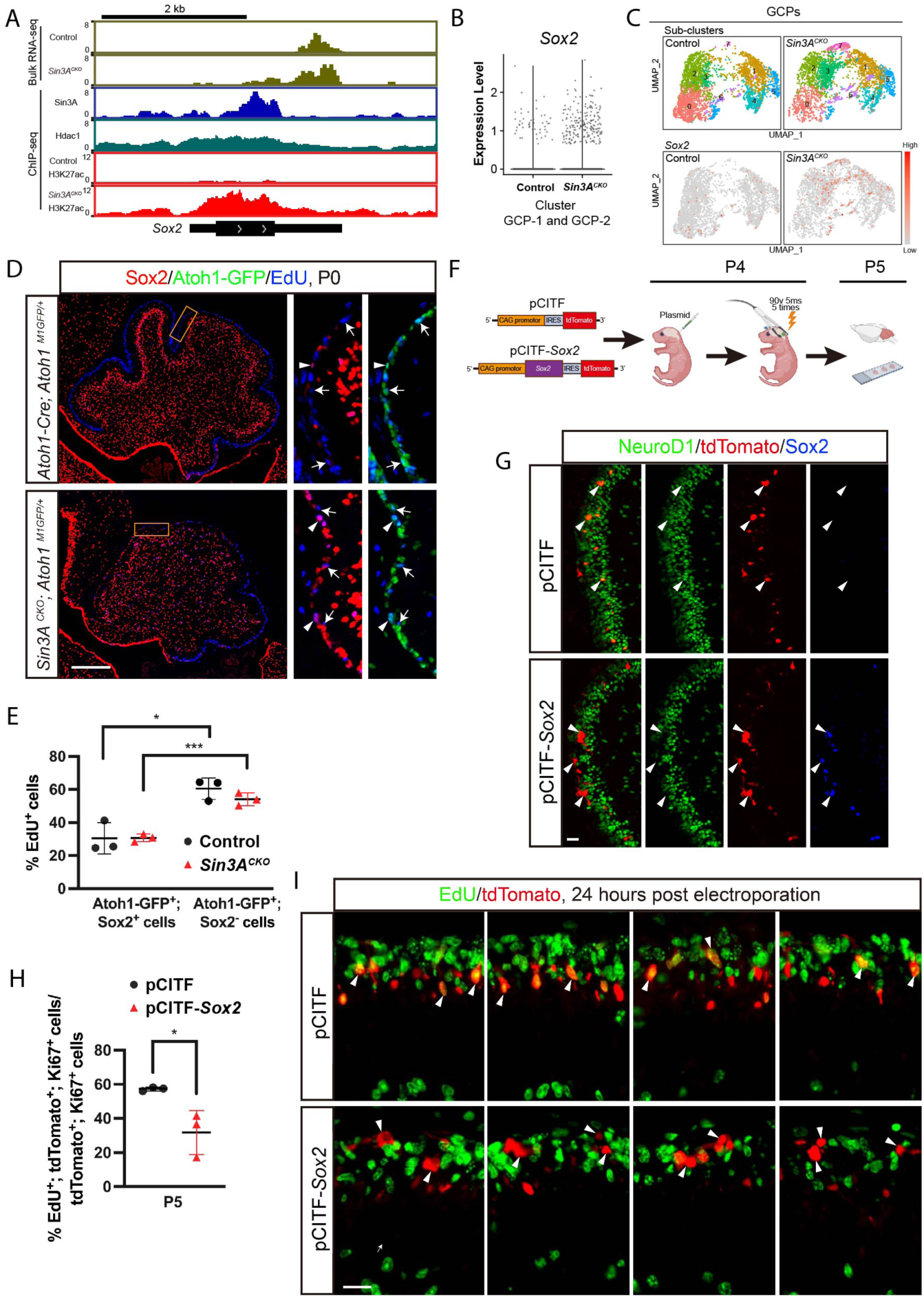
Sin3A represses *Sox2* expression to promote progression of Sox2^+^; Atoh1*^+^*progenitors. (A) Bulk RNA-seq tracks (reads per million values) showing the expression level of *Atoh1* in the GCPs of control and *Sin3A^CKO^*mice. ChIP-seq tracks (fold changes of ChIP-seq signal relative to IgG) depicting the intensity of Sin3A, Hdac1, and H3K27ac occupation at the *Sox2* locus in the dissociated cerebellar cells or isolated GCPs. The italicized text to the left of each track denotes the genotype of the mice used in ChIP-seq experiments; absence of genotype annotation indicates a wild-type or *Atoh1^M1GFP/+^*animal. (B) Violin plots showing the expression of *Sox2* in the GCP population of *Sin3A^CKO^* and control cerebellum. (C) Splitted UMAP displaying the *Sox2* expression in the sub-clusters of GCPs between *Sin3A^CKO^*and control. (D) Sox2, Atoh1-GFP, and EdU staining in the cerebellum of control and *Sin3A^CKO^* mice at P0. Mice were administrated with EdU 30 minutes before sacrifice. Arrowheads indicate cells positive for both Sox2 and EdU, and arrows indicate cells positive for both Atoh1-GFP and EdU. Scale bar indicates 250 μm. (E) Quantitative analysis showing the percentage of EdU^+^ cells relative to the Sox2^+^; Atoh1-GFP^+^ or Atoh1-GFP^+^ population in the EGL. Shown are mean ± SEM. Two-tailed t test: * p≤ 0.05, *** p ≤ 0.001. n= 3 mice per group. (F) Schematic representation of the strategy employed to overexpress *Sox2* using electroporation. (G) NeuroD1, tdTomato and Sox2 staining in the cerebellum of electroporated mice harvested at P5. Arrowheads indicate tdTomato^+^ cells in the EGL. Scale bar indicates 20 μm. (H) Quantitative analysis showing the percentage of EdU^+^ cells relative to the tdTomato^+^; Ki67^+^ population in the EGL. Shown are mean ± SEM. Two-tailed t test: * p≤ 0.05. n= 3 mice per group. (I) tdTomato and EdU staining in the cerebellum of electroporated mice at P5. Mice were administrated with EdU 30 minutes before sacrifice. Arrowheads indicate tdTomato^+^ cells in the EGL. Scale bar indicates 20 μm.

### Sin3A represses *Sox2* expression to promote progression of Sox2^+^; Atoh1*^+^* progenitors

Our transcriptome and ChIP-seq data have uncovered that *Sox2* is a direct target of the Sin3A/Hdac1 complex. This is supported by the elevated H3K27ac levels at Sox2 locus in *Sin3A^CKO^* GCPs (Fig. 5A). Moreover, analysis of scRNA-seq data at P6 indicate that *Sox2* is overexpressed in GCP-1 and GCP-2 clusters in the mutant cerebella (Fig. 5B and S3C). A detailed sub-clustering analysis of GCPs revealed that while *Sox2^+^* cells are rare in the GCPs of control cerebella, they are more abundant in *Sin3A^CKO^* GCPs, primarily distributed across multiple sub-clusters (Fig. 5C). The abundance of *Sox2^+^* cells increased nearly 5-fold in *Sin3A^CKO^* and constituted about 6.4% of the GCPs population as compared to 1.2% in the control.

At E13.5, both *Sin3A^CKO^* and control EGL exhibit a high proportion of Sox2^+^; Atoh1^+^ cells (Fig. 4A and 4E), as the process of exiting Sox2 expression has not yet commenced. By E16.5, the *Sin3A^CKO^*showed a significantly higher number of Sox2^+^; Atoh1^+^ cells compared to controls (Fig. 4B4), with their distribution extending further into the anterior EGL (Fig. 4B3). These cells represent 34.2% of the GCP population in the medial and anterior EGL of the *Sin3A^CKO^* cerebellum, which is 2.5 times greater than that observed in controls (Fig. 4E). However, the expression pattern of Sox2 remains similar to that in controls within both the RL and posterior EGL (Fig. 4B and S4B), consistent with the absence of Cre activity in those regions (Fig. 1B and S1). The increase in Sox2^+^; Atoh1^+^ cells is also evident at P0 and P7, accounting for 30.4% and 4.48% of Atoh1-GFP+ cells, respectively, compared to only 2.5% and 0.08% in controls (Fig. 4C3 to 4C4, 4D and 4E). Notably, these mutant Sox2^+^; Atoh1^+^ cells also displayed lower EdU labeling during the 30-minute EdU pulsing (Fig. 5D and 5E), although their S-phase duration did not show significant changes (Fig. S5A). This suggests a prolonged overall cell cycle for these Sox2^+^; Atoh1^+^ cells, thereby reducing the proportion of time spent in the S phase. Consequently, the overrepresentation of these slow-cycling cells in the EGL can lead to a reduced GCP pools, thereby decreasing the overall size of the GC lineage. This is corroborated by our observations that cell numbers in *Sin3A^CKO^*EGL were significantly lower than those in control from E16.5 to P7 (Fig. 4I), particularly at P0, where both the morphology and cell count in the anterior EGL remain at embryonic levels (Fig. 4C and 4I). Moreover, the previously observed cell apoptosis events did not occur within the Sox2^+^; Atoh1^+^ population (Fig. S5B). This indicates that despite a significant delay, these mutant cells eventually transform into Atoh1^+^ GCPs.

We next investigated whether Sin3A-mediated *Sox2* repression promotes the transition of Sox2^+^; Atoh1^+^ progenitors into Atoh1^+^ GCPs. To this end, we induced overexpression of Sox2 in the EGL of P4 mice along with tdTomato using *in vivo* electroporation (Fig. 5F). Twenty-four hours post-electroporation, tdTomato+ cells were observed within the cerebellar EGL. These cells co-expressed Sox2 but did not express NeuroD1, resembling Sox2^+^; Atoh1^+^ progenitors (Fig. 5G). In contrast, tdTomato+ cells in the control retained NeuroD1 expression (Fig. 5G). Furthermore, the proportion of EdU-labeled cells among those Sox2 overexpressing cells was reduced to approximately half that of the control (Fig. 5H and 5I). The result suggests that sustained Sox2 expression as observed in *Sin3A^CKO^* dampens cell cycle kinetics and impede GC lineage progression.

In addition to the Sox2^+^; Atoh1^+^ cells, we also observed another class of Sox2^+^ cells that are Atoh1-negative and located in the inner EGL (arrows in Fig. 4B1 to 4B4, and 4C1 to 4C4, 4D). These cells exhibited higher levels of Sox2 expression, similar to non-GC-lineage cells such as Bergmann glia and oligodendrocytes found in the deeper layers of the cerebellum. Their distribution pattern suggests that they may represent a class of Nestin-expressing progenitors (NEPs) that do not belong to the GC lineage, as previously reported.^62,63^ Although they have the potential to differentiate into GC cells upon injury or genetic defects,^63,64^ the absence of Atoh1 activity indicates that their *Sox2* expression remains unaffected by *Sin3A* conditional deletion. Indeed, lineage tracing using *Sin3A^CKO^; Ai9* mice confirmed that these cells were negative for tdTomato (Fig. S5C), indicating that they never expressed Atoh1 and do not belong to the GC lineage.

In summary, the transition of relatively quiescent progenitors from Sox2^+^; Atoh1^+^ to rapidly dividing Atoh1^+^ GCPs is a crucial driver for the rapid expansion of the GC lineage, a process supported by the epigenetic silencing of Sox2 expression by the Sin3A/Hdac1 complex.

### Sin3A-Hdac1 binds to *Atoh1* enhancers and represses *Atoh1* expression in GCPs

In addition to *Sox2*, we also identified *Atoh1* among 96 Sin3A target genes. *Atoh1* expression in the cerebellum requires the 1.5 kb autoregulatory enhancer located ∼3.4 kb 3’ of intronless *Atoh1* coding sequence (Fig. 6A).^65,66^ The 3’ regulatory region is composed of two enhancers (A and B) that are highly conserved across vertebrate species^67^ and required for Atoh1 function.^68^ The enhancer A contains a single degenerated E-box, whereas enhancer B contains an Atoh1 E Box Associated Motif (AtEAM) as reported previously (Fig. 6A).^26,34^ Analysis of Atoh1 ChIP-seq data set from P5 GCPs^26^ revealed that Atoh1 binds to not only the 3’ enhancer but also the 5’ region (Fig. 6A). The same regions are also occupied by Sin3A and Hdac1 and show extensive H3K27ac marks, especially at the 3’ enhancers A and B, in *Sin3A^CKO^* GCPs. Intriguingly, the occupancy of H3K27ac at the *Atoh1* enhancer in control GCPs is relatively low despite high chromatin accessibility (Fig. 6A). This observation suggests that Sin3A actively promotes deacetylation of H3K27 at the *Atoh1* enhancer in GCPs, potentially facilitating subsequent trimethylation of H3K27, a chromatin mark indicative of gene silencing (Fig. S6A).

**Figure 6.**
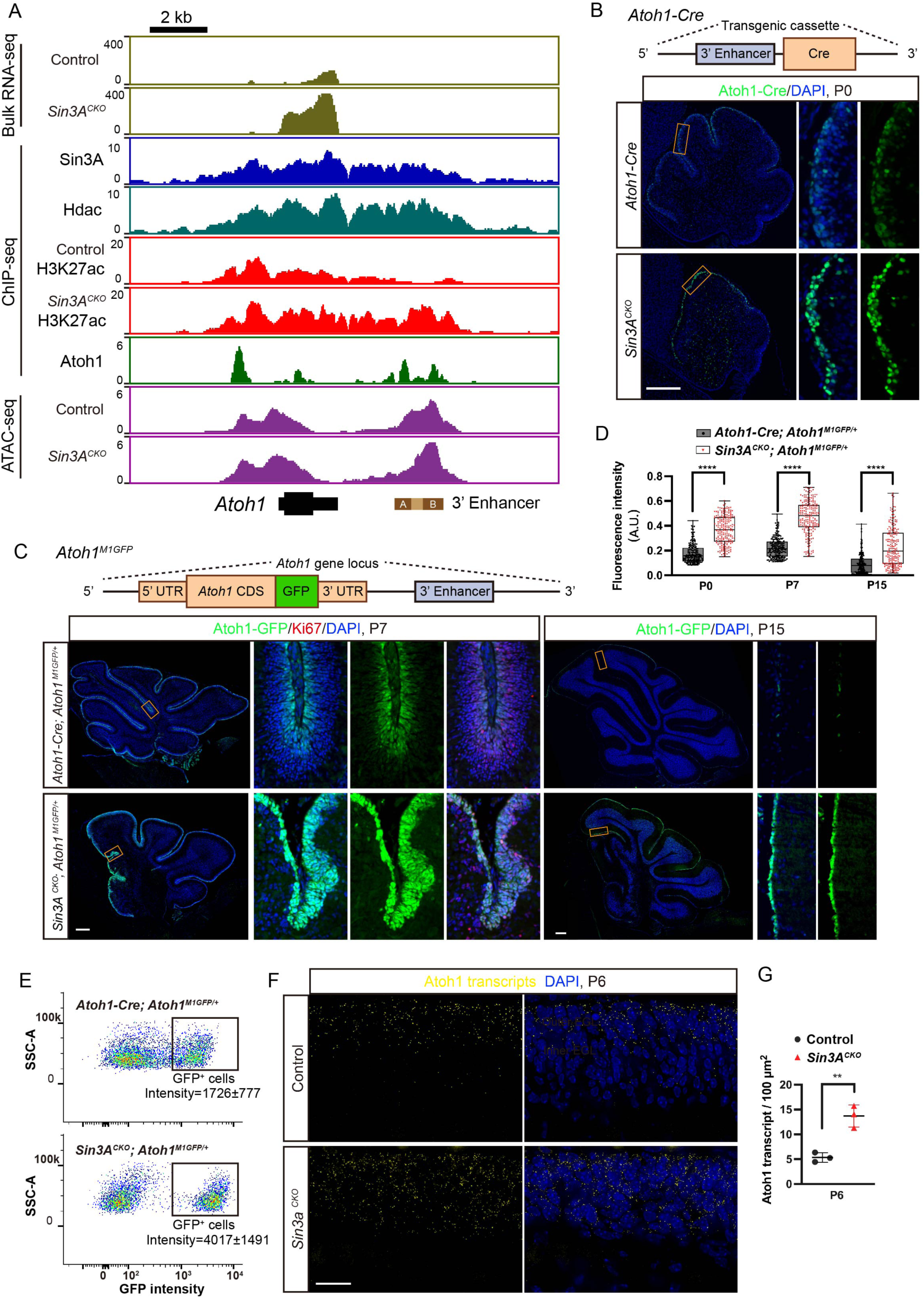
Sin3A-Hdac1 binds to *Atoh1* enhancers and represses *Atoh1* expression in GCPs. (A) Bulk RNA-seq tracks (reads per million values) showing the expression level of *Atoh1* in the GCPs of control and *Sin3A^CKO^*mice. ChIP-seq tracks (fold changes of ChIP-seq signal relative to IgG) depicting the intensity of Sin3A, Hdac1, and H3K27ac occupation at *Atoh1* locus in the dissociated cerebellar cells or isolated GCPs. The italicized text to the left of each track denotes the genotype of the mice used in ChIP-seq experiments; absence of genotype annotation indicates a wild-type or *Atoh1^M1GFP/+^*animal. Enhancers A and B are two critical regulatory elements located downstream of the Atoh1 coding region, identified through reported gene assay and cross-species conservation sequence analysis^65^. (B) Immunofluorescence staining for Atoh1-Cre at P0 in the cerebellum of control and *Sin3A^CKO^* mice. Nuclei were counterstained with DAPI. Scale bar indicates 250 μm. (C) Immunofluorescence staining for Atoh1-GFP and Ki67 at P7 and P15 in the cerebellum of control and *Sin3A^CKO^* mice. Nuclei were counterstained with DAPI. Scale bar indicates 250 μm. (D) Boxplot showing the quantification of the relative fluorescence intensity of GFP or Cre in EGL cells of control and *Sin3A^CKO^* mice. Two-tailed t test: **** p ≤ 0.0001. n= 200 cells per group. (E) Flow cytometry analysis of GCPs isolated from *Atoh1-Cre; Atoh1^M1GFP/+^* and *Sin3A^CKO^; Atoh1^M1GFP/+^* mice. Cells were dissociated from the anterior regions of the cerebellum in P7 mice and subsequently sorted based on endogenous GFP expression. (F) Single-molecule fluorescence in situ hybridization (smFISH) analysis showing the transcription levels of *Atoh1* in EGL of control and *Sin3A^CKO^* mice. Nuclei were counterstained with DAPI. Scale bar indicates 20 μm. (G) Quantification of the count of Atoh1 transcripts per 100 square micrometers. Shown are mean ± SEM. Two-tailed t test: ** p≤ 0.01. n= 3 mice per group.

The *Atoh1* repression by Sin3A was further validated by elevated expression of Cre and Atoh1-GFP from *Atoh1-cre* and *Atoh1 ^M1GFP^* alleles, respectively, in *Sin3A^CKO^* when compared to the control (Fig. 6B-6D). We observed ∼2 to 2.5-fold increase in Cre or Atoh1-GFP expression as determined by fluorescent intensity as early as P0 and continues throughout the presence of GCPs (Fig. 6B-6D). At P15, when the majority of GCPs expressing GFP have exited the cell cycle and migrated to IGL, mutant GCPs with elevated GFP expression persisted in the EGL (Fig. 6C). We also used FACS to quantitatively compare Atoh1-GFP expression between *Sin3A^CKO^*and control GCPs derived from the anterior region of the cerebella at P6. Consistent with immunohistochemistry data, we observed a greater proportion of cells with higher Atoh1-GFP intensity in the *Sin3A^CKO^* than the control (Fig. 6E). The mean Atoh1-GFP intensity was over 2-fold higher in the mutant, which is likely an underestimate due to the contamination of adjacent Sin3A^+^ cells during dissection. Indeed, RT-PCR data from these samples showed that the *Sin3A* transcript is still present at 30% level of the control (Fig. S6B). We also employed single molecule fluorescence in situ hybridization (smFISH) to spatially quantify *Atoh1* transcript levels. We found that the molecules of *Atoh1* transcript are significantly enriched in the EGL of *Sin3A^CKO^* mice (Fig. 6F and 6G). In control cerebellum, Atoh1 displayed a progressively diminishing distribution pattern from the outer to the inner EGL, with Atoh1 transcripts entirely absent in the innermost two cell layers. Conversely, the distribution of Atoh1 transcripts in *Sin3A^CKO^*cerebellum has extended to the innermost cells of EGL (Fig. 6F). Collectively, these studies support our model that Sin3A represses *Atoh1* expression by promoting deacetylation of H3K27ac at the *Atoh1* enhancer.

### NeuroD1 associate with the Sin3A/Hdac1 complex to repress Atoh1 expression

Due to its inability to directly bind DNA, the Sin3A/Hdac1 complex must recruit bona fide transcription factors to provide locus-specific gene regulations.^69,70^ To determine the factors associated with the Sin3A/Hdac1 complex in GCP, we conducted motif enrichment analysis on candidate genomic sites potentially regulated by the Sin3A/Hdac1 complex, utilizing sequence characteristics. We first identified genomic loci directly regulated by the Sin3A/Hdac1 complex, which are characterized by concurrent binding of both Sin3A and Hdac1 and exhibit differential levels of H3K27 acetylation upon depletion of Sin3A. A total of 3,439 genomic sites met these criteria (Fig. 7A). Mean profile plots and heatmaps of ChIP-seq signal illustrating the enrichment of H3K27ac revealed substantial hyperacetylation around these sites in the genome of *Sin3A^CKO^* (Fig. 7B). To predict which transcription factors mediate Sin3A/Hdac1 complex-induced deacetylation in these regions, we searched enriched motifs using Homer2.^71^ Notably, motifs corresponding to a group of b-HLH family members, including NeuroD1, Neurog2 and Atoh1, are highly enriched in the 3,439 sites (Fig. 7C and Supplementary table 2). Among these, NeuroD1 has shown to play an important role in the differentiation and survival of GCPs.^51,72–74^ NeuroD1 null mutants have reduced cerebellum size, elevated cell death and Atoh1 expression,^51,75^ a phenotype strikingly similar to that observed in *Sin3A^CKO^*. Moreover, the expression of NeuroD1 in the EGL exhibits a gradual increase from the outer to the inner layers, which is inversely correlated with the decreasing trend of Atoh1 (Fig. 7D and S6C). Notably, this negative correlation between NeuroD1 and Atoh1 is dependent on Sin3A as indicated by the persistent Atoh1 expression in majority of NeuroD1^+^ cells lacking Sin3A function (Fig. 7D and S6C). Particularly in regions where granule cell precursors abnormally cluster together, many cells co-express high levels of both Atoh1 and NeuroD1 (Fig. 7D). Therefore, NeuroD1 appears to be a prime DNA-binding candidate that associate with the Sin3A/Hdac1 complex to repress *Atoh1* expression.

**Figure 7.**
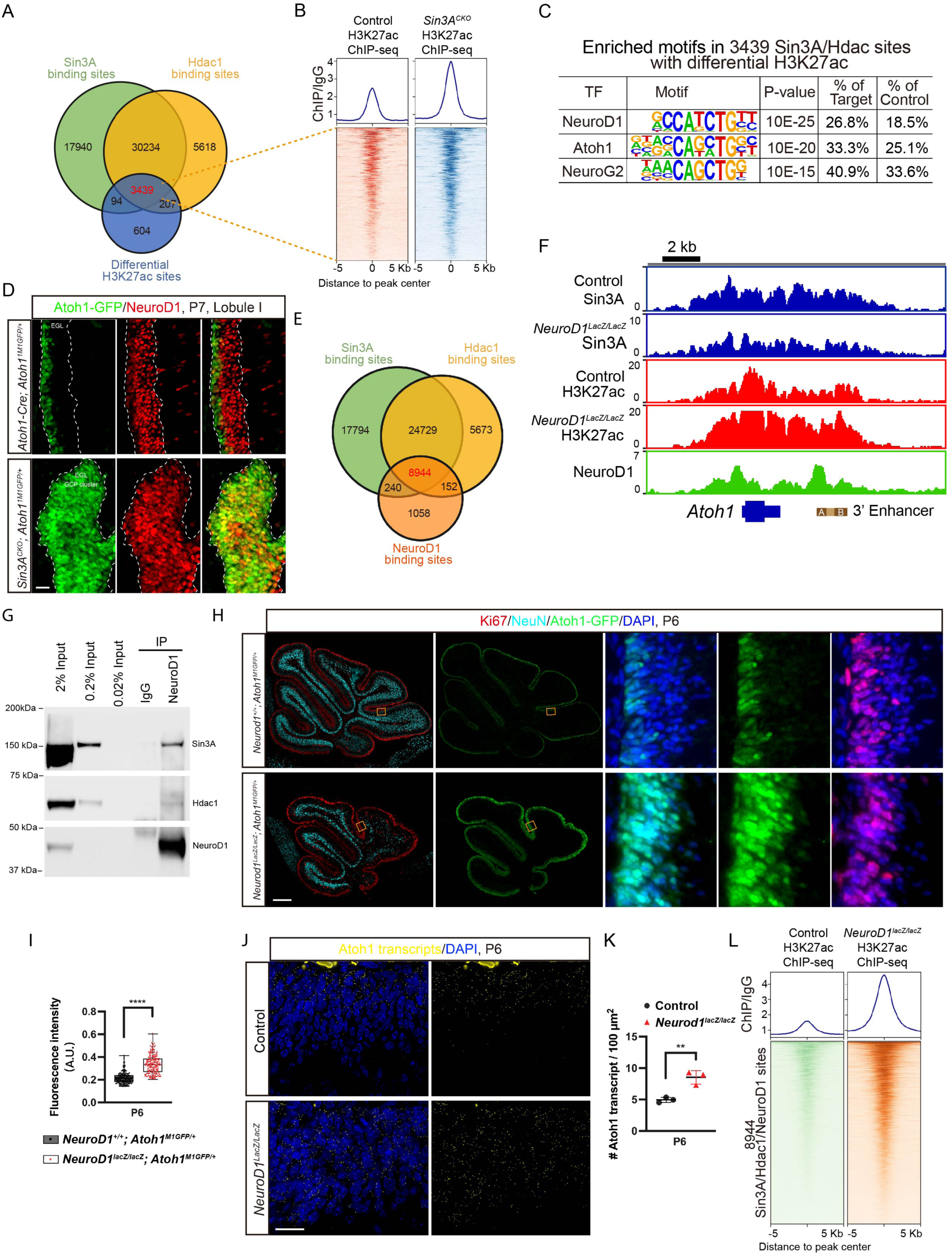
NeuroD1 associate with the Sin3A/Hdac1 complex to repress Atoh1 expression. (A) Venn diagram illustrating the identification of genomic loci directly regulated by the Sin3A/Hdac1 complex. (B) Heatmap and average intensity plots illustrating the H3K27ac ChIP-seq signal profiles surrounding 3439 genomic loci directly regulated by the Sin3A/Hdac1 complex in isolated GCP from both control and *Sin3A^CKO^* mice. (C) Selected motifs exhibit significant enrichment at genomic loci that are directly regulated by the Sin3A/Hdac1 complex. (D) Immunofluorescence staining for Atoh1-GFP and NeuroD1 at P6 in the cerebellum of control and *Sin3A^CKO^*mice. Scale bar indicates 250 μm. (E) Venn diagram illustrating the overlaps of Sin3A, Hdac1 and NeuroD1 binding sites in the genome of wild type cerebellum. (F) ChIP-seq tracks (fold changes of ChIP-seq signal relative to IgG) depicting the intensity of Sin3A, Hdac1, NeuroD1, and H3K27ac occupation at *Atoh1* locus in the GCP of control, *Sin3A^CKO^*, and *NeuroD1 ^LacZ/LacZ^* mice. The italicized text to the left of each track denotes the genotype of the mice used in ChIP-seq experiments, absence of genotype annotation indicates a wild-type animal. (G) Immunoblots for NeuroD1, Sin3A and Hdac1 binding in P7 cerebellum of wild-type mice. Lane 1-3: cell lysate of P7 cerebellum. Lane 4: Co-immunoprecipitation of rabbit IgG in cell lysate of P7 cerebellum. Lane 5: Co-immunoprecipitation of NeuroD1 in cell lysate of P7 cerebellum. (H) Immunofluorescence staining for Atoh1-GFP, NeuN, and Ki67 at P6 in the cerebellum of control and *NeuroD1 ^LacZ/LacZ^* mice. Nuclei were counterstained with DAPI. Scale bar indicates 20 μm. (I) Boxplot showing the quantification of the relative fluorescence intensity of GFP in EGL cells of control and *NeuroD1^lacZ/lacZ^* mice. Two-tailed t test: **** p ≤ 0.0001. n= 200 cells per group. (J) smFISH analysis showing the transcription levels of Atoh1 in the EGL of wild type and *NeuroD1^lacZ/lacZ^*mice. Nuclei were counterstained with DAPI. Scale bar indicates 20 μm. (K) Quantification of the count of Atoh1 transcripts per 100 square micrometers in the EGL of *wild type* and *NeuroD1^lacZ/lacZ^* mice. Shown are mean ± SEM. Two-tailed t test: ** p≤ 0.01. n= 3 mice per group. (L) Heatmap and average intensity plots of the H3K27ac ChIP-seq signals across 10 Kb from the center of Sin3A/Hdac1/NeuroD1 binding sites throughout the genome from isolated GCPs of control and *NeuroD1^lacZ/lacZ^*mice.

We next tested whether NeuroD1 binds to the Atoh1 locus by performing ChIP-seq experiment using the chromatin extracted from cerebella of P6 wildtype mice. Our analysis of ChIP-seq data have identified a total of 10,394 NeuroD1 binding sites, of which 86.04% overlap with Sin3A/Hdac1 co-binding sites (Fig. 7E). These loci bound by Sin3A, Hdac1, and NeuroD1 represent 17.6% of all genomic Sin3A binding sites. There is a modest enrichment of NeuroD1 occupancy around Sin3A binding sites (Fig. S6D), indicating a strong association between the DNA-binding patterns of NeuroD1 and Sin3A. Notably, at Atoh1 locus, NeuroD1 bound to the promoter and the 3’ enhancer, regions that overlapped with the binding sites of both Sin3A and Hdac1 (Fig. 7F). Consistent with the notion that NeuroD1 associates with Sin3A/Hdac1 as a complex, we found a direct physical interaction between NeuroD1, Sin3A, and Hdac1 in the developing cerebellum using co-immunoprecipitation (Fig. 7G).

To further investigate the functional correlation between NeuroD1 and Sin3A in transcriptional regulation of *Atoh1*, we employed *NeuroD1^lacZ^*knock-in mouse line in which the entire coding region of NeuroD1 has been replaced by a β-galactosidase reporter gene. This modification resulted in the loss of NeuroD1 expression ^75^. Consistent with previous studies,^51,75^ *NeuroD1^lacz/lacZ^*mice exhibited a reduction in cerebellar volume and a decrease in the number of lobules (Fig. 7H). Similar to *Sin3A^CKO^* mice, there is an increase in the thickness of the proliferative cell layer and expansion of Atoh1-GFP that spans almost the entire depth of the EGL. Moreover, relative expression levels of Atoh1-GFP were 1.56-fold higher in the cerebellum of *NeuroD1^lacz/lacZ^*mice than controls (Fig. 7H and 6I). We also performed smFISH analysis and found that *Atoh1* transcripts were similarly expanded and elevated in *NeuroD1^lacz/lacZ^* cerebella (Fig. 7J and 7K).

We reasoned that if NeuroD1 acts as a component of the Sin3/Hdac1 complex to repress *Atoh1* gene expression, its absence would likely alter epigenetic states of *Atoh1* locus as observed in Sin3A deficiency. Indeed, deficiency in NeuroD1 resulted in a genome-wide increase in H3K27 acetylation levels (Fig. 7L), consistent with the hyperacetylation observed in the Sin3A-deficient model (Fig. S6E). Notably, there was a 4.6-fold elevation in H3K27ac levels at the *Atoh1* locus (Fig. 7F), which bears a striking resemblance to that identified in *Sin3A^CKO^*.

To further support NeuroD1 is an essential component of the Sin3A/Hdac1 complex that represses *Atoh1* expression, we conducted Sin3A ChIP-seq using chromatin from NeuroD1-null cerebella to determine potential changes in the binding profile of Sin3A. The results indicate that the binding peak of Sin3A at the *Atoh1* locus is reduced by 2.8-fold in the absence of NeuroD1 (Fig. 7F), highlighting the importance of NeuroD1 for recruiting the Sin3A/Hdac1 complex to the *Atoh1* enhancer.

Collectively, the results indicate that the Sin3A/Hdac1 complex targets the *Atoh1* gene locus through its interaction with NeuroD1, thereby inhibiting its transcriptional activity.

## Discussion

The cerebellar EGL comprises a diverse population of lineally related cells with distinct molecular signatures and differentiation potentials. These cells organize a lineage hierarchy essential for GC development, in which slow-cycling Sox2⁺ progenitors sit at the apex and transiently localize to the EGL’s outer edge during embryonic and early postnatal stages.^9^ Our findings reveal that the majority of Sox2^+^ cells also co-express Atoh1, affirming their role as transitory progenitors for rapidly dividing GCPs. These SOX2^+^; ATOH1^+^ cells exhibit cross-species conservation, as they have been identified in both the EGL of the 17 post coitus week (PCW) human cerebellum and in ATOH1 lineage cells derived from human pluripotent stem cells (hPSCs).^76^ This observation correlates with analogous Sox2^+^; Atoh1^+^ transient cells identified in hair cell lineages, where Sox2 blockade initiates the hair cell differentiation program.^77^ Furthermore, we demonstrate that during embryonic development, Sox2^+^; Atoh1^+^ cells in the EGL are not a rare cell type. Instead, they represent a transitional cell type necessary for the development from Sox2^+^ ECPs to Atoh1^+^ GCPs. At E13.5, these cells constitute over 90% of the EGL population. After then their proportion rapidly decreases, rendering them a rare cell type by birth.

As a core regulatory factor for maintaining stem cell characteristics,^21^ Sox2 plays a crucial role in controlling the cell cycle. Gain- and loss-of-function experiments have elegantly demonstrated that the downregulation of Sox2 is sufficient to accelerate cell cycle progression, prompting neural stem cells to exit their quiescent state and rapidly proliferate, generating more new neurons; conversely, overexpression of Sox2 reverses this process.^23,78^ Mechanistically, the effect of Sox2 in maintaining cellular quiescence may be achieved through its inhibition of genes associated with the cell cycle and mitosis.^79^ Once neural stem cells commit to a neuronal fate, the expression of Sox2 is rapidly downregulated.^24,80^ Our results demonstrate that in the vast majority of post-commitment GC lineage cells, Sox2 must also be timely suppressed. Failure to do so will result in these cells remaining in a relatively primitive state with slow cycling instead of transitioning into rapidly proliferating GCPs, thereby affecting GCP population expansion. However, before its expression ceases, the role of Sox2 in GC lineage cells remains unclear. Although our electroporation experiments have demonstrated that ectopic expression of Sox2 is sufficient to revert Atoh1^+^ GCPs into more primitive quiescent transitional cells, conditional knockout of Sox2 using hGFAP-Cre—which becomes active in the EGL starting from E14.5—does not result in any noticeable phenotypic changes.^81,82^ This phenomenon, where a loss-of-function mutation shows no phenotype while a gain-of-function mutation does, suggests that the function of Sox2 may be compensated by other members of the SoxB1 family. For instance, Sox1, which is also expressed in Sox2^+^; Atoh1^+^ cells, shows slightly increased expression in Sin3A^CKO^ samples according to bulk RNA-seq data.

As the apex of the GC lineage hierarchy, the developmental origin of Sox2^+^ progenitors in the EGL was not been thoroughly described. Based on their distribution characteristics, gene expression profiles, and lineage tracing results, we propose that they represent a transitional state between Sox2^+^ ECPs and Atoh1^+^ GCPs. Unlike the recently questioned transitional cells between neural stem cells (NSCs) and GCPs in the human cerebellum,^83,84^ the Sox2^+^; Atoh1^+^ cells we identified exhibit comparable unique molecular identifiers (UMIs) and gene counts to other cell clusters (Fig. S4E). Furthermore, this cluster of transitional cells is consistently observed across several independent datasets (Fig. S4H). Therefore, it can be ruled out as a technical artifact resulting from low-quality single-cell RNA-seq data.

Unlike Sox2^+^; Atoh1^+^ progenitors, rapidly proliferating GCPs undergo four to six rounds of cell division before exiting the cell cycle and differentiating into GCs.^85^ Atoh1 maintains GCPs in the precursor state through an autoregulatory loop reliant on Atoh1 binding to E-box motifs within its 3’ enhancers.^65,68^ Perturbations in Atoh1 levels have shown to either impede or promote the transition from GCPs to GCs.^27,29,31^ Notably, the transcriptional repressor Hic1 can suppress Atoh1 expression when overexpressed in dissociated GCPs.^86^ However, Hic1 does not appear to bind Atoh1 enhancers until late differentiation stages, suggesting additional factors are necessary to initiate the GCP-to-GC transition. We propose that NeuroD1 fulfills this critical role by recruiting the Sin3A/Hdac1 complex to Atoh1 enhancers to suppress expression. This is supported by several observations: (1) NeuroD1 physically associates with Sin3A and Hdac1; (2) NeuroD1 binds directly to Atoh1 enhancers; (3) Sin3A binding at the Atoh1 locus is significantly reduced without NeuroD1; (4) H3K27ac levels at the Atoh1 locus are sharply elevated in NeuroD1’s absence; (5) Atoh1 transcripts persist and accumulate without NeuroD1. Thus, the GCP-to-GC transition occurs via dynamic shifts in Atoh1 expression from positive autoregulation to a negative feedback loop where Atoh1 induces NeuroD1 expression,^26^ which, in turn, suppresses Atoh1 expression.

Like in the cerebellum, NeuroD1 plays a crucial role in differentiating various neurons across brain regions, including the dentate gyrus, hippocampus, cerebral cortex, and olfactory system.^75,87–89^ Additionally, NeuroD1 can reprogram different cell types into neuronal phenotypes neurons.^90–92^ This reprogramming capability is linked to NeuroD1’s ability to bind closed chromatin regions and convert them into an open state, marked by the loss of repressive H3K27me3 and gain of active H3K27ac marks.^90,93^ Accordingly, NeuroD1 is widely recognized as a transcriptional activator. NeuroD1’s repressor role is often indirect, activating or displacing repressors on chromatin.^94^ In microglia-to-neuron transdifferentiation, NeuroD1 directly activates transcriptional repressors Scrt1 and Meis2, which are involved in microglial identity suppression.^90^ In murine embryonic stem cells, NeuroD1 displaces enhancer-bound Tbx3, a neuronal fate repressor, promoting neuronal gene expression.^93^ Our finding that NeuroD1 can associate with the Sin3A/Hdac1 complex to suppress Atoh1 expression introduces a novel paradigm where Neurod1 recruits co-factors with diverse histone modification activities to regulate gene transcription in a context dependent manner.

We also noted that the removal of Neurod1 did not entirely eliminate the Sin3A binding capability at the Atoh1 locus, suggesting additional transcriptional repressors recruit the Sin3A/Hdac1 complex to suppress Atoh1 expression. This aligns with observations that GCP differentiation isn’t completely abolished without Neurod1, as some GCs are still generated.^73^ While other Neurod1 family members aren’t expressed in the EGL, distant neural-specific bHLH factors Nhlh1 and Nhlh2 are present in the EGL and expected to play roles in GCP differentiation.^95,96^ Future studies are necessary to elucidate their regulatory relationships with Atoh1.

Negative regulation of Atoh1 at the protein level also promotes GCP differentiation. Impaired Atoh1 protein degradation leads to similar defects as Sin3A mutants, with arrested proliferative states and failure to differentiate.^31,97^ This suggests precise multilayered control over Atoh1 expression blockade. The transcriptional suppression we’ve observed occurs at the most upstream level of gene expression, which may account for the higher proliferation in *Sin3A^CKO^* models.

Aside from increased Atoh1 expression, *Neurod1^KO^* and *Sin3A^CKO^* models share many phenotypes, including hypoplastic cerebellum, thickened EGL, persistent GCP proliferation, and increased cell death.^51^ However, a noteworthy difference is that apoptotic cells lacking NeuroD1 predominantly appear in the inner EGL, where differentiation begins, whereas in *Sin3A^CKO^* models, NeuroD1 expression persists and apoptotic cells distribute throughout the EGL. Cell death in NeuroD1-deficient cells is linked to a lack of downstream neurotrophic factors.^73^ These observations indicate distinct mechanisms driving GCP apoptosis in Sin3A mutants. In mouse embryonic stem cells, Sin3A is essential for preventing or repairing DSBs, and its absence results in DNA damage and p53-independent cell cycle arrest and apoptosis.^98^ The presence of DNA double-strand breaks as indicated by γ-H2AX in *Sin3A^CKO^* GCPs suggests a similar requirement of Sin3A to inhibit DNA damage-induced apoptosis.

Shh-subtype medulloblastoma (SHH-MB) is thought to arise from GC lineage precursors^99,100^ and relies on Atoh1 for tumor initiation.^30,86^ Similar to the development of GC lineage, SHH-MB also follows a hierarchical pattern.^9,101,102^ Heterogeneous cell subpopulations within the tumor tissue recapitulate different developmental stages of GC lineage cells.^101^ Our discovery that Sin3A’s stage-specific functions in GC lineage progression suggests a more intricate role for Sin3A in SHH-MB. Early in the lineage, Sin3A promotes the expansion of rapidly proliferating Atoh1^+^ GCPs and potentially influences the corresponding MB cell subpopulation; later, it restrains Atoh1 activity to drive GCP/MB differentiation. These district functions are likely mediated by specific transcription factor partner, including NeuroD1 identified in this study. NeuroD1 is highly expressed in differentiated medulloblastoma cells and has been shown to inhibit tumor growth when overexpressed in animal models.^74,103^ Our finding that the Sin3A/Hdac1/NeuroD1 complex suppresses Atoh1 expression provides mechanistic insight into NeuroD1’s function in medulloblastoma.

The cerebellar hypoplasia phenotype, identified in the *Sin3A^CKO^* mouse model, may provide an explanation for the cerebellar atrophy and cerebellar ataxia symptoms observed in some cases of Witteveen-Kolk syndrome.^48,104^ Although we did not observe significant cerebellar developmental defects in heterozygous mice, this discrepancy might be attributed to differences between mice and humans regarding tolerance to haploinsufficiency.^105^ Additionally, the widely reported ventriculomegaly phenotype in human patients suggests potential cortical hypoplasia.^42,104,106^ This implies that Sin3A might play a crucial regulatory role during the development of other neurons in the brain.

## Resource availability

### Lead contact

Further information and requests for regents and resources should be directed to and will be fulfilled by the lead contact, Chin Chiang.

### Materials availability

All materials and reagents generated in this study are available on request from the lead contact.

### Data and code availability

- ChIP-seq raw and processed data are available through the Gene Expression Omnibus under accession GEO: GSE309720. To access them, please use reviewer token: wdknaucgbbotxqb. scRNA-seq and bulk RNA-seq raw data are available through the the European Nucleotide Archive under accession ENA: PRJEB98573 and PRJEB98570, respectively. ATAC-seq raw data are available through ENA: PRJEB98574. Accession numbers are listed in the key resources table.
- This paper does not report original code.
- Any additional information required to reanalyze the data reported in this work paper is available from the lead contact upon request.

## Acknowledgements

We thank Mark Magnuson, M.D. for sharing *Neurod1^lacZ^* mice. We also thank Guoqiang Gu, Ph.D, for advice on using insulin to extend the lifespan of *Neurod1^lacZ/lacZ^* mutants. This study was supported by grants to C.C. from the Vanderbilt-Ingram Cancer Center Support Grant P30 CA068485 and the National Institutes of Health NS 131495.

## Author contributions

L.C. and C.C. designed the experiments and wrote the paper. L.C. and A.R. performed all the experiments. L.C. and C.C. analyzed data. G.D. provided the mice with *Sin3A* floxed allele.

## Declaration of interests

The authors declare no conflict of interests.

## STAR★Methods

### Key resources table

| REAGENT or RESOURCE | SOURCE | IDENTIFIER |
| --- | --- | --- |
| **Antibodies** |  |  |
| GFP | Aves Labs | Cat# GFP-1020,<br>RRID:AB_10000240 |
| Cleaved Caspase-3 | Cell Signaling Technology | Cat# 9661,<br>RRID:AB_2341188 |
| Ki67 | Thermo Fisher Scientific | Cat# 14-5698-82,<br>RRID:AB_10854564 |
| Cdkn1b (p27) | Cell Signaling Technology | Cat# 3698,<br>RRID:AB_2077832 |
| NeuroD1 | Cell Signaling Technology | Cat# 62953,<br>RRID:AB_2077832 |
| Sin3A | Novus Biologicals | Cat# NB600-1263,<br>RRID:AB_2187631 |
| Hdac1 | Cell Signaling Technology | Cat# 34589,<br>RRID:AB_2756821 |
| Sox2 | Thermo Fisher Scientific | Cat# 740013T,<br>RRID:AB_3093719 |
| Ccnd1 | Cell Signaling Technology | Cat# 55506,<br>RRID:AB_2827374 |
| Cleaved Caspase-3 (Asp175) | Cell Signaling Technology | Cat# 9664,<br>RRID:AB_2070042 |
| Cre Recombinase | Cell Signaling Technology | Cat# 15036,<br>RRID:AB_2798694 |
| Acetyl-Histone H3 (Lys27) | Cell Signaling Technology | Cat# 8173,<br>RRID:AB_10949503 |
| NeuN | Cell Signaling Technology | Cat# 24307,<br>RRID:AB_2651140 |
| Goat anti - mouse Alexa Fluor 488 | Thermo Fisher Scientific | Cat# A-11017,<br>RRID:AB_2534084 |
| Goat anti - rabbit Alexa Fluor 488 | Thermo Fisher Scientific | Cat# A-11008,<br>RRID:AB_143165 |
| Goat anti - mouse Alexa Fluor 568 | Thermo Fisher Scientific | Cat# A-11004,<br>RRID:AB_2534072 |
| Goat anti - rabbit Alexa Fluor 647 | Thermo Fisher Scientific | Cat# A-21244,<br>RRID:AB_2535812 |
| Goat anti - chicken Alexa Fluor 488 | Thermo Fisher Scientific | Cat# A-11039,<br>RRID:AB_2534096 |
| Donkey anti - goat Alexa Fluor 647 | Thermo Fisher Scientific | Cat# A-21447,<br>RRID:AB_2535864 |
| Donkey anti - rabbit Alexa Fluor 488 | Thermo Fisher Scientific | Cat# A-21206,<br>RRID:AB_2535792 |
| **Bacterial strains** |  |  |
| NEB® 5-alpha Competent *E. coli* (High Efficiency) | New England Biolabs | Cat# C2987I |
| **Reagent** | **Source** | **Identifier** |

| **Chemicals, Peptides, and Recombinant Proteins** |  |  |
| --- | --- | --- |
| AMPure XP SPRI Reagent | Beckman Coulter | A63880 |
| Nextera DNA CD Indexes | Illumina | 20018707 |
| Dynabeads™ Protein A for Immunoprecipitation | Thermo Fisher Scientific | 10001D |
| Dynabeads™ Protein G for Immunoprecipitation | Thermo Fisher Scientific | 10003D |
| 5-ethyl-2'-deoxyuridine (EdU) | Click Chemistry Tools | 1149-500 |
| KAPA2G Fast HotStart Genotyping Mix | Millipore | 2GFHSGKB |
| NaCl | Sigma-Aldrich | S7653 |
| KCl | Sigma-Aldrich | P9333 |
| MgCl<sub>2</sub>·6H<sub>2</sub>O | Research Products International | 7791-18-6 |
| KH<sub>2</sub>PO<sub>4</sub> | Sigma-Aldrich | P0662 |
| NaHCO<sub>3</sub> | Sigma-Aldrich | S5761 |
| CaCl<sub>2</sub>·2H<sub>2</sub>O | Sigma-Aldrich | 223506 |
| D-glucose | Sigma-Aldrich | G8270 |
| Disuccinimidyl glutarate | Thermo Fisher Scientific | 20593 |
| HEPES | Sigma-Aldrich | H3375 |
| EGTA | Research Products International | 67-42-5 |
| Protease from Bacillus licheniformis | Sigma-Aldrich | P5380 |
| Deoxyribonuclease I from bovine pancreas | Sigma-Aldrich | DN25 |
| Insulin glargine (Lantus) | Sanofi | 3238193 |
| Tn5 | Illumina | 20034197 |
| AMPure XP Beads | Beckman Coulter | A63880 |
| Liberase TL | Roche | 540102000 |
| **Software and algorithms** |  |  |
| FlowJo (V.10.4) | Three Star | https://www.flowjo.com/ |
| ImageJ (V.1.52a) | NIH | https://imagej.nih.gov/ij/ |
| Prism 10 (V.10.2.3) | GraphPad Software | https://www.graphpad.com/scientific-software/prism/ |
| Adobe Illustrator 2021 | Adobe | https://www.adobe.com/ |
| ZEN (V.2.6) | Carl Zeiss | https://www.zeiss.com/microscopy/de/produkte/software/zeiss-zen.html |
| R (V.4.1.0) | The R Project for Statistical Computing | https://www.r-project.org/ |
| Seurat (v.4.4.0) |  | https://satijalab.org/seurat/ |
| Deseq 2 (v.1.38.3) |  | https://bioconductor.org/packages/3.16/bioc/html/DESeq2.html |
| Homer (v.4.11) |  | http://homer.ucsd.edu/homer/index.html |
| BioRender | BioRender | https://app.biorender.com/ |
| Cellprofiler (V.4.2.8) | Broad Institute | https://cellprofiler.org/ |
| **Critical commercial assays** |  |  |
| RNeasy Micro Kit | Qiagen | Cat# 74004 |
| Click-iT™ Plus TUNEL Assay Kits for In Situ Apoptosis Detection | Thermo Fisher Scientific | Cat# C10619 |
| SMART-Seq® v4 Ultra® Low Input RNA Kit for Sequencing | Takarabio | Cat# 634888 |
| NEBNext® Ultra™ II DNA Library Prep Kit for Illumina® | New England Biolabs | Cat# E7645S |
| **Deposited data** |  |  |
| Control rep1, GCP, H3K27ac (ChIP) | This paper | Accession #GSM9277963 |
| Control rep2, GCP, H3K27ac (ChIP) | This paper | Accession #GSM9277964 |
| *Sin3A<sup>CKO</sup>* rep1, GCP, H3K27ac (ChIP) | This paper | Accession #GSM9277965 |
| *Sin3A<sup>CKO</sup>* rep2, GCP, H3K27ac (ChIP) | This paper | Accession #GSM9277966 |
| Control, GCP, IgG (ChIP) | This paper | Accession #GSM9277967 |
| *Sin3A<sup>CKO</sup>*, GCP, IgG (ChIP) | This paper | Accession #GSM9277968 |
| *NeuroD1<sup>lacZ/lacZ</sup>* rep1, GCP, H3K27ac (ChIP) | This paper | Accession #GSM9277969 |
| *NeuroD1<sup>lacZ/lacZ</sup>* rep2, GCP, H3K27ac (ChIP) | This paper | Accession #GSM9277970 |
| NeuroD1 control rep1, GCP, H3K27ac (ChIP) | This paper | Accession #GSM9277971 |
| NeuroD1 control rep2, GCP, H3K27ac (ChIP) | This paper | Accession #GSM9277972 |
| Wildtype rep1, Cerebellum, Sin3A (ChIP) | This paper | Accession #GSM9277973 |
| Wildtype rep2, Cerebellum, Sin3A (ChIP) | This paper | Accession #GSM9277974 |
| *NeuroD1<sup>lacZ/lacZ</sup>* rep1, Cerebellum, Sin3A (ChIP) | This paper | Accession #GSM9277975 |
| *NeuroD1<sup>lacZ/lacZ</sup>* rep2, Cerebellum, Sin3A (ChIP) | This paper | Accession #GSM9277976 |
| Wildtype rep1, Cerebellum, Hdac1 (ChIP) | This paper | Accession #GSM9277977 |
| Wildtype rep2, Cerebellum, Hdac1 (ChIP) | This paper | Accession #GSM9277978 |
| Wildtype, Cerebellum, (Input) | This paper | Accession #GSM9277979 |
| Wildtype, Cerebellum, IgG (ChIP) | This paper | Accession #GSM9277980 |
| Wildtype rep1, Cerebellum, NeuroD1 (ChIP) | This paper | Accession #GSM9277981 |
| Wildtype rep2, Cerebellum, NeuroD1 (ChIP) | This paper | Accession #GSM9277982 |
| Wildtype rep1, GCP (RNA-seq) | This paper | Accession #ERR15665402 |
| Wildtype rep2, GCP (RNA-seq) | This paper | Accession #ERR15665403 |
| *Sin3A<sup>CKO</sup>* rep1, GCP (RNA-seq) | This paper | Accession #ERR15665404 |
| *Sin3A<sup>CKO</sup>* rep2, GCP (RNA-seq) | This paper | Accession #ERR15665405 |
| Cerebellum_2XWildtype+2X*Sin3A<sup>CKO</sup>*_4-plex (scRNA-seq) | This paper | Accession #ERR15665326 |
| GCP_Wildtype (ATAC-seq) | This paper | Accession # ERR15665653 |
| GCP_*Sin3A<sup>CKO</sup>* (ATAC-seq) | This paper | Accession # ERR15665654 |
| Wildtype, Cerebellum, P0, H3K27me<sup>3</sup> (ChIP-seq) | Pal, et al.<sup>107</sup> | Accession # GSM587804 |
| Wildtype, Cerebellum, P5, H3K27me<sup>3</sup> (ChIP-seq) | Pal, et al.<sup>107</sup> | Accession # GSM587805 |
| Wildtype, Cerebellum, P15, H3K27me<sup>3</sup> (ChIP-seq) | Pal, et al.<sup>107</sup> | Accession # GSM587806 |
| Wildtype, Cerebellum, Adult, H3K27me<sup>3</sup> (ChIP-seq) | Pal, et al.<sup>107</sup> | Accession # GSM587807 |
| *Atoh1*<sup>FLAG/FLAG</sup>, Cerebellum, P5, Atoh1 (ChIP-seq) | Klisch, et al.<sup>26</sup> | Accession # GSM549935 |
| Wildtype, Cerebellum, E13.5 (scRNA-seq) | Wizeman et al.<sup>108</sup> | Accession #GSM3399102 |
| Wildtype, Cerebellum, E14 (scRNA-seq) | Khoury-Farah et al.<sup>109</sup> | Accession #GSM5393629 |
| Wildtype, Hindbrain, E12 (scRNA-seq) | Vladoiu, et al.<sup>61</sup> | Accession #GSM3318000 |
| Wildtype, Hindbrain, E14 (scRNA-seq) | Vladoiu, et al.<sup>61</sup> | Accession #GSM3318001 |
| Wildtype, Hindbrain, E16 (scRNA-seq) | Vladoiu, et al.<sup>61</sup> | Accession #GSM3318002 |
| **Recombinant DNA** |  |  |
| pCITF-Sox2 | In house | Subcloned from Addgene #61404 and # #13462 |
| **Oligonucleotides** |  |  |
| Stellaris FISH Probes, Custom Assay with Quasar® 670 Dye | LGC Biosearch Technologies | Cat# SMF-1065-5 |
| *Sin3a*<sup>Flox</sup> | In house<sup>49</sup> |  |
| *Atoh1*<sup>M1GFP</sup> | The Jackson Laboratory.<sup>59</sup> | Stock No. 013593<br>RRID:IMSR_JAX:013593 |
| *Atoh1*<sup>Cre</sup> | The Jackson Laboratory.<sup>110</sup> | Stock No. 011104<br>RRID:IMSR_JAX:011104 |
| *Neurod1*<sup>lacZ</sup> | Gift from Mark Magnuson, M.D.<sup>75</sup> | RRID:MGI:6220689 |
| *Ai9* | The Jackson Laboratory.<sup>54</sup> | Stock No. 007909<br>RRID:IMSR_JAX:007909 |

## Experimental model and study participant details

### Mice

All experimental procedures adhered to animal care and biosafety protocols approved by the Vanderbilt University Division of Animal Care, in compliance with NIH guidelines. Histological analyses were conducted on specimens ranging from E16.5 to P40. RNA-seq and ChIP-seq experiments used P6 to P7 mice. The following mouse lines were used in this study:

1. *Sin3A^Flox^*, LoxP sites were introduced to flank exon 4 of the *Sin3A* gene. The functional knockout of *Sin3A* is based on the cell-specific activity of Cre recombinase.^49^
2. *Atoh1-Cre*, a transgenic mouse line with a random insertion of a cassette inclued 1.5 kb Atoh1 3’ enhancer fragment driving the expression of Cre recombinase.^110^
3. *Atoh1^M1GFP^*, a knock-in mouse line with a fused EGFP to the C-terminus of Atoh1 protein.^59^
4. *Ai9* (*R26R-tdTomato)*, A CAG loxP-stop-loxP-tdTomato cassette was inserted into the ROSA26 locus, enabling the expression of tdTomato under the control of the CAG promoter in a Cre recombinase-dependent manner.^54^
5. *NeuroD1^lacZ^*, *NeuroD1 knock-out mouse* line in which the entire coding region of NeuroD1 has been replaced by a β-galactosidase reporter gene.^75^

## Method details

### Mice treatments

For the analysis of proliferation and differentiation, pups at P0, P5, and P7 were administered an intraperitoneal injection of 50 mg/kg EdU (Click Chemistry Tools). Their brains were collected either 30 minutes or 48 hours after the injection and subsequently prepared for paraffin embedding. For *NeuroD1^lacZ/lacZ^* pups, subcutaneous injections of 0.1 U insulin glargine (Lantus, Sanofi) were administered twice daily to alleviate symptoms of diabetes.

### Electroporation

The pCITF or pCITF-Sox2 plasmid was dissolved in Tris-EDTA buffer at a concentration of 3 µg/µL, with the addition of 0.15% Fast Green dye. Following hypothermic anesthesia on ice, P4 mice were injected with 3 µL of the plasmid DNA solution into the cerebellum using a microsyringe with a 34-gauge needle (Hamilton). Subsequently, electroporation was performed using an ECM830 electroporator (BTX) equipped with tweezer-type electrodes; the negative electrode was placed at the occipital regions and the positive electrode under the mandible. Five pulses of 90 V were delivered, each separated by a 50 ms interval. After electroporation, mice were placed on a heating pad maintained at 37°C until full recovery from anesthesia was observed.

### Histological analysis

Harvested cerebellum were fixed in 4% paraformaldehyde (PFA) for 48 hours. Subsequently, they were processed sagittally into either 5 μm paraffin sections or 15 μm cryosections. Sections from adult mice were stained with hematoxylin and eosin solution. Immunofluorescence staining was performed according to previously described protocols.^111^ Primary antibodies used in this study are listed below: GFP (1:1000, Aves Labs, cat#: GFP-1020), Ki67 (1:1000, Thermo Fisher Scientific, cat#: 14-5698-82), Cdkn1b (1:500, Cell Signaling Technology, cat#: 3698), NeuroD1 (1:1000, Cell Signaling Technology, cat#: 62953), Sin3A (1:200, Novus Biologicals, cat#: NB600-1263), Sox2 (1:500, Thermo Fisher Scientific, cat#: 740013T), Ccnd1 (1:300, Cell Signaling Technology, cat#: 55506), Cleaved Caspase-3 Asp175 (1:200, Cell Signaling Technology, cat#: 9664), Cre Recombinase (1:500, Cell Signaling Technology, cat#: 15036), NeuN (1:500, Cell Signaling Technology, cat#: 24307). Image acquisition was carried out using a Leica DMi8 microscope imaging system or a Zeiss LSM880 confocal microscope.

### Single-molecule fluorescence in situ hybridization (smFISH)

The probe library targeting Atoh1 transcripts is composed of 48 specific probes, each 20 bp in length, designed using the Stellaris Online Probe Designer (https://www.biosearchtech.com/products/rna-fish/), and synthesized with Quasar^®^ 670 Dye by Biosearch Technologies. SmFISH was performed according to previously described protocols.^112^ In brief, cryosections of 6 μm thickness were mounted onto poly-L-lysine-coated cover slides and postfixed with a 3.7% formaldehyde/PBS solution for 15 minutes. Subsequently, the sections were permeabilized with 70% ethanol at 4°C overnight and equilibrated in wash buffer containing 30% formamide for six hours. Hybridization was then carried out overnight at 30°C using a solution containing fluorescent probes and with 30% formamide at a concentration of 0.1 ng/μl. Excess probes were removed by washing twice with wash buffer, followed by nuclear counterstaining with DAPI. Imaging was performed using a Leica DMi8 fluorescence microscopy system equipped with a 60X 1.4NA objective lens. Transcript molecule quantification was carried out utilizing the ImageJ software package.

### Cell sorting

After the removal of the meninges from the cerebellar tissue, it was sectioned into small pieces approximately 0.5 mm in size. Subsequently, these fragments were digested with a cold protease cocktail solution, which contained 10 mg/ml protease from bacillus licheniformis (Sigma-Aldrich) and 2.25 mg/ml deoxyribonuclease I from bovine pancreas (Sigma-Aldrich), for 15 minutes in a water bath maintained at 6℃, with gently pipetting to mechanically dissociate the tissues every five minutes during this period. After dissociation, the digestion enzyme mixtures were passed through a 40μm filter (Miltenyi Biotech) and washed twice by 1XPBS solution. The BD FACS Aria III cell sorter was used to purify GCPs based on the intensity of Atoh1-GFP fluorescence at the single-cell level.

### co-IP

Fresh cerebellar tissue was ground into a powder in liquid nitrogen, followed by the adding of RIPA buffer and incubation on ice for 20 minutes to lyse the cells. The lysate containing total soluble proteins was obtained after ultrasonication and subsequent centrifugation to collect the supernatant. Following the quantification of protein concentrations via the bicinchoninic acid (BCA) assay, lysates containing 10 ug protein were incubated with Dynabeads Protein A (Thermo Fisher Scientific) conjugated to anti-NeuroD1 antibody (1:100, Cell Signaling Technology, cat#: 62953) at 4°C for a period of 2 hours. In control experiments, nonspecific rabbit IgG was used. Subsequently, magnetic beads were washed five times with wash buffer (10 mM Tis-HCl; 150 mM NaCl; 0.5 mM EDTA; 0.05% NP40) to remove unbound proteins. Afterthen, the proteins bound to the beads were eluted using a 100 mM lysine solution at pH 2.8 and analyzed by SDS-PAGE electrophoresis. The PVDF membrane containing proteins was subjected to immunoblot analysis using anti-Sin3A (1:1000, Novus Biologicals, cat#: NB600-1263), anti-Hdac1 (1:1000, Cell Signaling Technology, cat#: 34589), and anti-NeuroD1 (1:1000, Cell Signaling Technology, cat#: 62953) antibodies.

### ATAC-seq

A total of 50,000 Atoh1-GFP-positive GCPs were isolated from the anterior cerebellum of both *Sin3A^CKO^* and control mice to generate ATAC-seq libraries. Cells were lysed in a lysis buffer (10 mM Tris-HCl, 10 mM NaCl, 3 mM MgCl_2_, 0.1% IGEPAL CA-630), and nuclei were collected by centrifugation. Subsequently, chromatin-accessible regions were labeled using the transposase activity of Tn5 (Illumina). The DNA was then purified with AMPure XP Beads (Beckman Coulter). Finally, the ChIP DNA was amplified to create a sequencing library using Illumina Nextera DNA CD Indexes (Illumina). All libraries underwent multiplex paired-end 150 bp sequencing on an Illumina NovaSeq 6000 at Vanderbilt Technologies for Advanced Genomics.

### RNA-seq

A total of 1,000 Atoh1-GFP-positive GCPs were isolated from the anterior cerebellum of both *Sin3A^CKO^* and Control mice to generated RNA-seq sequencing libraries. Cellular RNA was extracted and cDNA was generated using the SMART-Seq v4 Ultra Low Input RNA Kit for Sequencing (Takarabio) according to manufacturer’s instructions. NEBNext Ultra II DNA Library Prep Kit for Illumina (New England Biolabs) was utilized to construct libraries from cDNA. All libraries are sequenced and performed at multiplex Paired-End 150 bp on an Illumina NovaSeq 600 at Vanderbilt Technologies for Advanced Genomics.

### ChIP-seq

ChIP-seq was followed by the ChIPmentation protocol^113^. In brief, purified cells were cross-linked with 1% formaldehyde/PBS solution at room temperature for 15 minutes before quenching with 250 mM glycine. For co-repressor (Sin3A and Hdac1) ChIP assays, an additional protein cross-linking step using 2mM disuccinimidyl glutarate (Thermo Fisher Scientific) was incorporated. Chromatin was subsequently fragmented to the size of 200∼1000bp by sonication for 21 seconds using a Diagenode One device (Diagenode). Soluble chromatin was then obtained and incubated overnight at 4°C with the target antibody. Afterward, Dynabeads Protein A/G (Thermo Fisher Scientific) were added for two hours to capture the antibody-chromatin complexes. Unbound chromatin was removed through multiple washes following a low-salt buffer (20 mM Tris-HCl, 500 mM NaCl, 2 mM EDTA, 1% Triton X-100, 0.1% SDS), a high-salt buffter (20 mM Tris-HCl, 150 mM NaCl, 2 mM EDTA, 1%Triton X-100, 0.1%SDS), a LiCl buffer (10 mM Tris-HCl, 250 mM LiCl, 1 mM EDTA, 1% IGEPAL CA-630, 0.5% DOC), and a 10mM Tris buffer sequence to ensure specificity of binding to the beads. Subsequently, an on-bead tagmentation step were performed using transposase Tn5 (Illumina) to facilitate the ligation of sequencing adapters to the DNA. The DNA was then eluted from the beads and purified using AMPure XP Beads (Beckman Coulter). Finally, the ChIP DNA was amplified to generate a sequencing library with Illumina Nextera DNA CD Indexes (Illumina). All libraries are sequenced and performed at multiplex Paired-End 150 bp on a Illumina NovaSeq 600 at Vanderbilt Technologies for Advanced Genomics. Primary antibodies used in this study are listed below: NeuroD1 (1:100, Cell Signaling Technology, cat#: 62953), Sin3A (1:100, Novus Biologicals, cat#: NB600-1263), Hdac1 (1:50, Cell Signaling Technology, cat#: 34589), Acetyl-Histone H3 Lys27 (1:100, Cell Signaling Technology, cat#: 8173).

### ChIP-seq and RNA-seq data analysis

For RNA-seq data, the FASTQ files were initially processed using Trim Galore to remove sequencing adapters, followed by alignment of the sequences to the Mus musculus mm10 genome with HISAT2. Subsequently, Samtools was employed to convert the mapping results into BAM format and generate corresponding indices. Quantification of read counts for all genes was performed using HTSeq-count, resulting in a matrix correlating samples with gene count numbers. Two biological replicates were examined. Differential expression analysis was conducted utilizing DESeq2, adopting a threshold of p < 0.01 and fold change > 2 as criteria for selecting differentially expressed genes.

For ChIP-seq data, the FASTQ files were initially processed using Cutadapt to remove sequencing adapter sequences, followed by alignment of the sequences to the mouse mm10 genome with Bowtie2. The aligned results were then converted into BAM files using Samtools and filtered to exclude reads aligning to mitochondrial DNA and regions on the blacklist. Additionally, PCR duplicates were removed utilizing Picard’s MarkDuplicates tool. Subsequently, peaks were called in comparison with IgG ChIP-seq controls using MACS2. For Sin3A, Hdac1, and NeuroD1 ChIP-seq analyses, two biological replicates were examined. Reproducible peaks were identified using the Irreproducible Discovery Rate method to establish a peak set for subsequent analysis. After obtaining genomic coordinates for these peaks, they were annotated within corresponding genomic regions via Homer’s annotatePeaks function. For comparing levels of H3K27ac across different genotypes of *Sin3A*, two biological replicates were examined, and DiffBind was employed for combination and differential analysis between samples, adopting a threshold of p < 0.01 and a fold change greater than 2 as criteria for selecting significantly different acetylation peaks.

### scRNA-seq

After dissecting lobules I to VII of the mouse cerebellum, the tissue was fixed and dissociated into a single-cell suspension following the 10X Tissue Fixation & Dissociation protocol (GC000553). Briefly, approximately 25 mg of cerebellar tissue was fixed in 1 ml of Fixation Buffer at 4°C for 20 hours. Subsequently, it was digested in a Dissociation Solution containing 0.2 mg/ml ml Liberase TL (Roche) at 37°C for 20 minutes. The dissociated tissue was then passed through a 30 µm filter to obtain a single-cell suspension. Single-cell transcriptomic profiling was performed with the 10x Genomics Chromium GEM-X Flex Gene Expression assay (Protocol G000691, Rev. A). Approximately 300,000 cells per probe barcode (minimum 25,000 cells/barcode) were hybridized with Chromium Mouse Transcriptome Probe Set v1.1.1(19,403 genes targeted by 55,479 probes) at 42 °C for 2 h in a thermomixer with heated lid. Following hybridization, unbound probes were removed via washes. Washed cell suspensions were loaded into the Chromium X controller along with GEM-X Flex chips and barcoded gel beads. Gel Beads-in-Emulsion (GEM) formation encapsulated single cells, barcoded beads, reverse transcription and ligation reagents. Within each GEM, adjacent left/right probe pairs hybridized to target transcripts and were ligated, incorporating cell- and molecule-specific barcodes (UMIs). GEMs were recovered, emulsions broken, and pooled cDNA amplified via PCR. Libraries were purified, quantified with Qubit (Thermo Fisher), and fragment sizes evaluated on an Agilent Bioanalyzer. Sequencing was performed on an Illumina NovaSeq 6000 with paired-end reads (R1: 28 bp cell barcode/UMI; R2: 90 bp probe insert), targeting 50,000– 100,000 read pairs per cell.

### scRNA-seq data analysis

Raw data were processed using Cell Ranger (v7.1.0) with alignment to mouse reference genome mm10. Gene counts per cell were generated based on UMI-corrected, ligated probe sequences and summarized in digital expression matrices. Only transcripts corresponding to targeted coding regions were captured due to probe-based specificity. Subsequently, quality control of the cells was performed based on gene count and UMI count, with doublets removed using scDblFinder 1.18.0. Normalization, dimension reduction, and clustering analysis were conducted using Seurat 4.4.0. Differential cell abundance analysis was carried out with miloR 2.0.0, while pseudo time trajectory analysis utilized CellRank 2.

### Quantification and statistical analysis

NIH Image J software was used to measure the area (μm^2^) for regions of interest (ROI) and for the acquisition of cell counting. For each stage, three midsagittal sections (5 μm thick) of each cerebellum were used for quantitative analysis. For quantification of cells in the EGL and IGL, the entire laminate was used as ROIs. All quantitative data were analyzed using Prism 10 (GraphPad) for statistical analysis and graphic presentations. Unpaired student’s t-test was used to compare the statistical difference between control and *Sin3A^CKO^*animals.

## Supplemental information

Document S1. Figures S1–S6

Document S2. Table S1–S3

Supplemental video

## Notes

### Competing Interest Statement

The authors have declared no competing interest.

