## Supplemental figures for "Hierarchical regulation of cerebellar neurogenesis by Sin3A-mediated gene repression"

#### Supplementary Figure legends

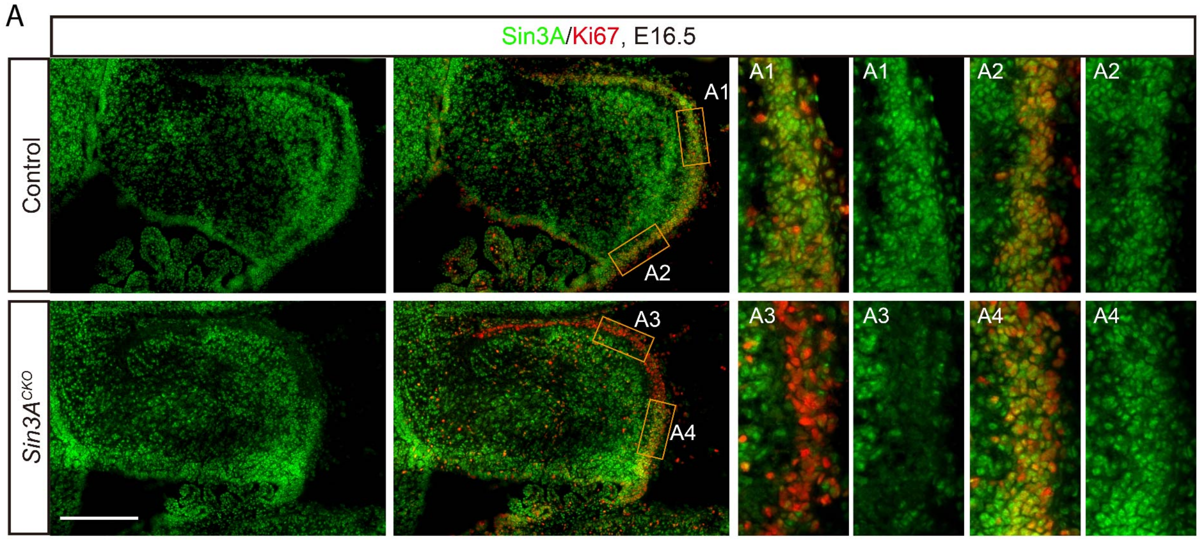


**Figure S1. Additional images of Sin3A expression in the embryonic EGL. Related to Figure 1.**

(A) Immunofluorescence staining for Ki67 and Sin3A at E16.5 in the cerebellum of control and *Sin3A*$^{CKO}$ mice. Panels A1 and A3 showing the anterior regions of EGL, while A2 and A4 showing the posterior regions. Scale bar indicates 250 µm.

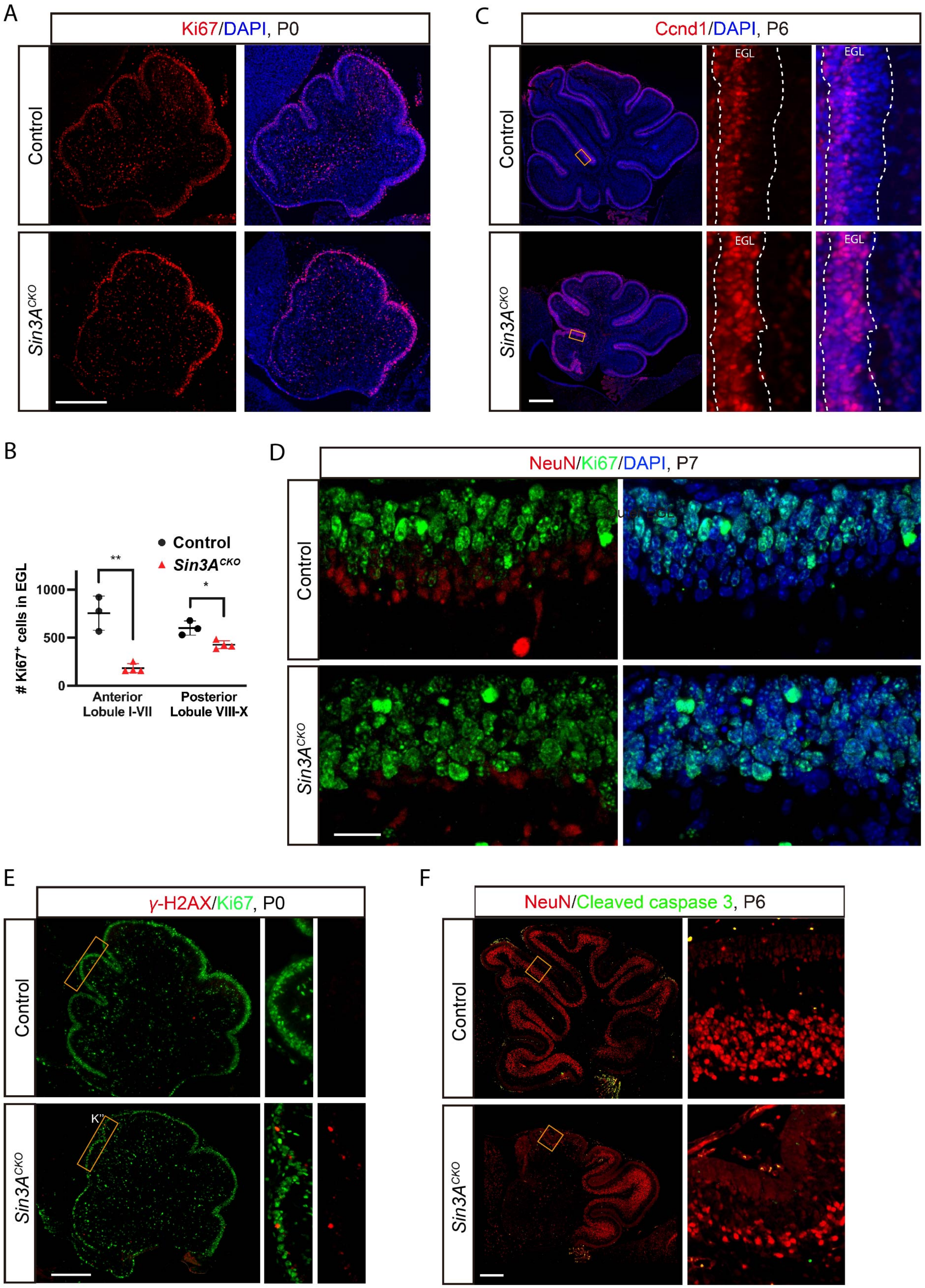

A
Ki67/DAPI, P0
Control
Sin3A^CKO
B
### Ki67+ cells in EGL
Control
Sin3A^CKO
**
*
1000
500
0
Anterior Lobule I-VII
Posterior Lobule VIII-X
C
Ccnd1/DAPI, P6
EGL
D
NeuN/Ki67/DAPI, P7
E
γ-H2AX/Ki67, P0
K''
F
NeuN/Cleaved caspase 3, P6

G

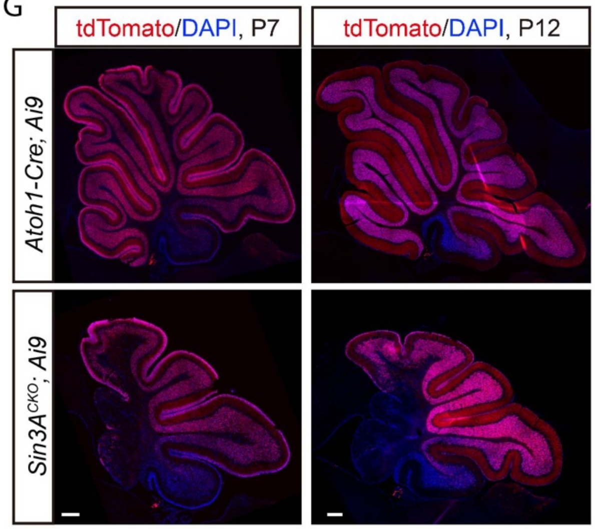


**Figure S2. Additional images showing cellular changes in the cerebellum of *Sin3A*$^{CKO}$ mice. Related to Figure 2.**

(A) Immunofluorescence staining for Ki67 at P0 in the cerebellum of control and *Sin3A*$^{CKO}$ mice. Nuclei were counterstained with DAPI. Scale bar indicates 250 μm.

(B) Quantification of the Ki67$^{+}$ cells in the EGL. Shown are mean ± SEM. Two-tailed t test: * p≤ 0.05, ** p≤ 0.01. n= 3 or 4 mice per group.

(C). Immunofluorescence staining for Ccnd1 at P6 in the cerebellum of control and *Sin3A*$^{CKO}$ mice. Nuclei were counterstained with DAPI. Scale bar indicates 250 μm.

(D). Immunofluorescence staining for Ki67 and NeuN at P7 in the cerebellum of control and *Sin3A*$^{CKO}$ mice. Nuclei were counterstained with DAPI. Scale bar indicates 20 μm.

(E). Immunofluorescence staining for Ki67 and γ-H2AX at P0 in the cerebellum of control and *Sin3A*$^{CKO}$ mice. Scale bar indicates 250 μm.

(F). Immunofluorescence staining for CP3 and NeuN at P6 in the cerebellum of control and *Sin3A*$^{CKO}$ mice. Scale bar indicates 250 μm.

(G). Immunofluorescence staining for tdTomato at P7 and P12 in the cerebellum of *Atoh1-Cre; Ai9* and *Sin3A*$^{CKO}$*; Ai9* mice. Nuclei were counterstained with DAPI. Scale bar indicates 250 μm.

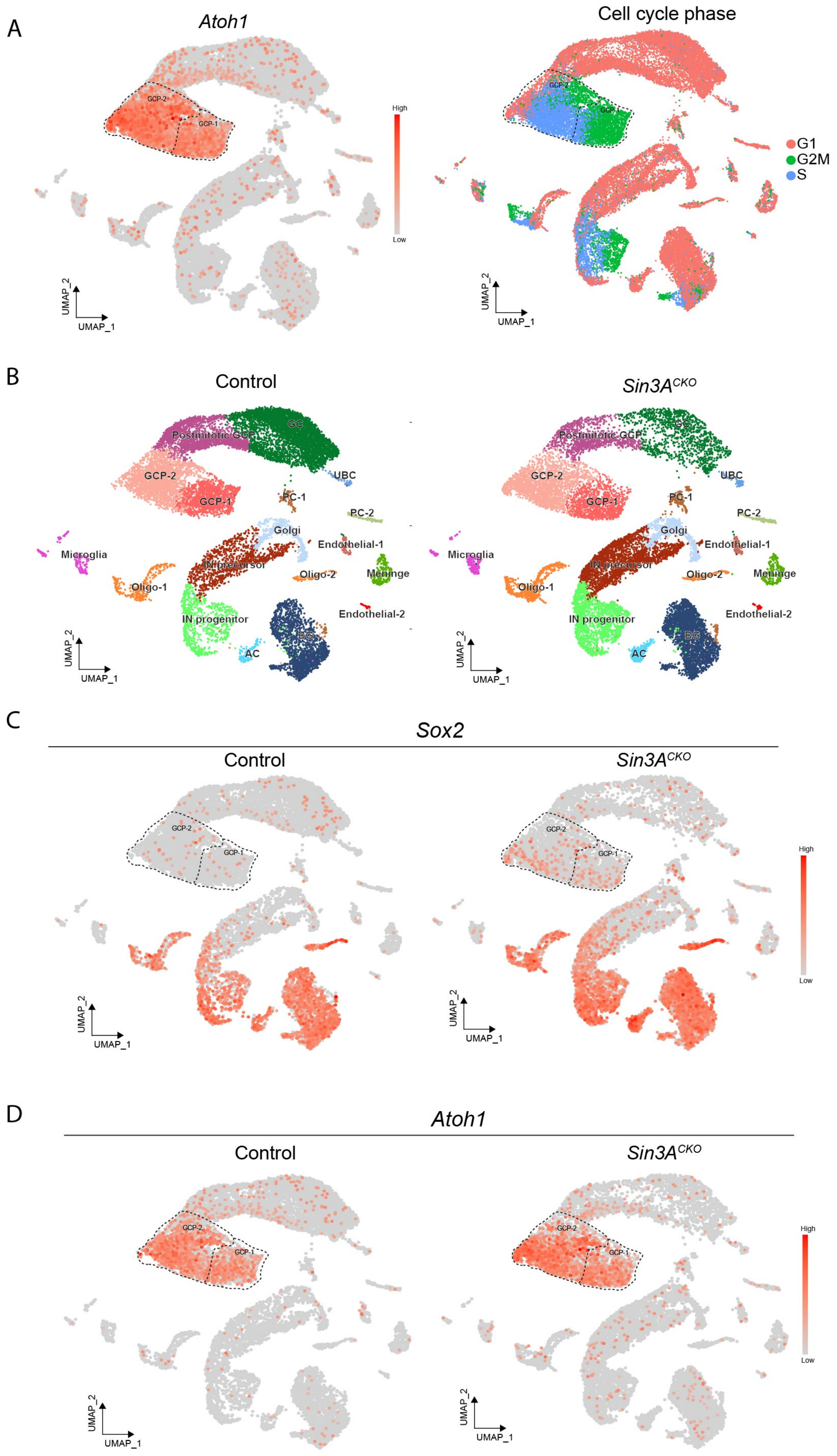
A
Atoh1
GCP-2
GCP-1
High
Low
UMAP_2
UMAP_1
Cell cycle phase
G1
G2M
S
B
Control
Sin3A^CKO
Postmitotic GCP
GC
GCP-2
GCP-1
UBC
PC-1
PC-2
Golgi
Endothelial-1
Microglia
IN precursor
Oligo-2
Meninge
Oligo-1
Endothelial-2
IN progenitor
BG
AC
C
Sox2
Control
Sin3A^CKO
D
Atoh1
Control
Sin3A^CKO

**Figure S3. Additional images showing cellular changes in the cerebellum of *Sin3A*$^{CKO}$ mice. Related to Figure 3, 4, and 6.**

(A) UMAP visualization of the *Atoh1* expression and cell cycle phase assignment.

(B) Splitted UMAP visualization of the cell clusters of cerebellum between *Sin3A*$^{CKO}$ and control.

(C) Splitted UMAP visualization of the *Sox2* expression between *Sin3A*$^{CKO}$ and control.

(D) Splitted UMAP visualization of the *Atoh1* expression between *Sin3A*$^{CKO}$ and control.

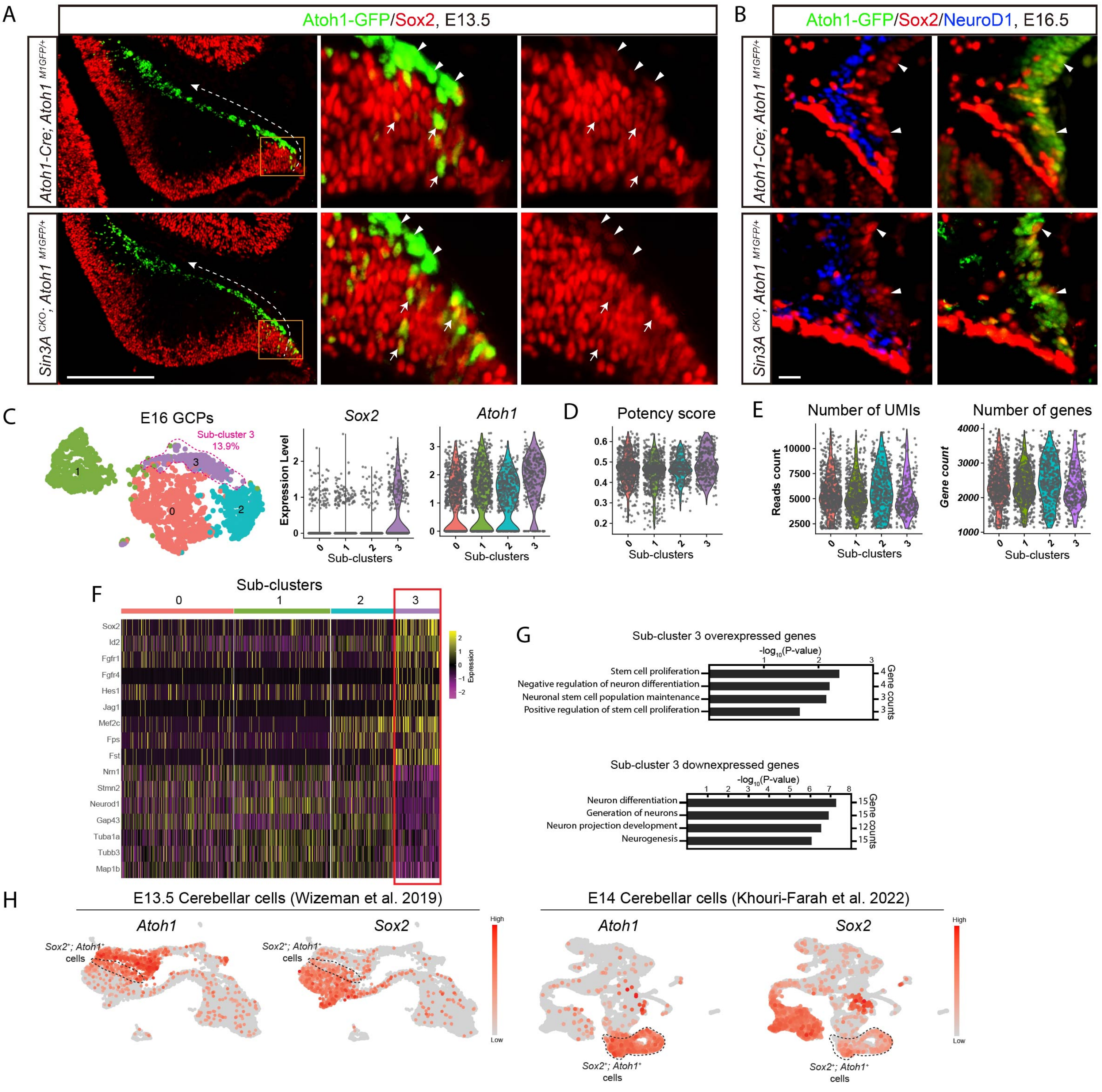
A
Atoh1-GFP/Sox2, E13.5
Atoh1-Cre; Atoh1 M1GFP/+
Sin3A CKO; Atoh1 M1GFP/+
B
Atoh1-GFP/Sox2/NeuroD1, E16.5
Atoh1-Cre; Atoh1 M1GFP/+
Sin3A CKO; Atoh1 M1GFP/+
C
E16 GCPs
Sub-cluster 3
13.9%
Sox2
Atoh1
Expression Level
Sub-clusters
D
Potency score
Sub-clusters
E
Number of UMIs
Reads count
Number of genes
Gene count
Sub-clusters
F
Sub-clusters
Sox2
Id2
Fgfr1
Fgfr4
Hes1
Jag1
Mef2c
Fps
Fst
Nrn1
Stmn2
Neurod1
Gap43
Tuba1a
Tubb3
Map1b
Expression
G
Sub-cluster 3 overexpressed genes
-log10(P-value)
Stem cell proliferation
Negative regulation of neuron differentiation
Neuronal stem cell population maintenance
Positive regulation of stem cell proliferation
Gene counts
Sub-cluster 3 downexpressed genes
-log10(P-value)
Neuron differentiation
Generation of neurons
Neuron projection development
Neurogenesis
Gene counts
H
E13.5 Cerebellar cells (Wizeman et al. 2019)
Atoh1
Sox2
Sox2+; Atoh1+ cells
High
Low
E14 Cerebellar cells (Khouri-Farah et al. 2022)
Atoh1
Sox2
Sox2+; Atoh1+ cells
High
Low

**Figure S4. Additional images showing Sox2$^+$; Atoh1$^+$ cells in the cerebellum. Related to Figure 4 and 5.**

(A) Immunofluorescence staining for GFP and Sox2 at E13.5 in the RL of control and *Sin3A*$^{CKO}$ mice. The dashed lines with arrows in the left panel indicate the direction of GC lineage migration. Arrows indicate Sox2$^{high}$; Atoh$^+$ cells in RL, and arrowheads indicate Sox2$^{low}$; Atoh$^+$ cells in uRL. Scale bar indicates 250 μm.

(B) Immunofluorescence staining for GFP, Sox2 and NeuroD1 at E16.5 in the RL of control and *Sin3A*$^{CKO}$ mice. Arrowheads indicate Sox2$^+$; Atoh$^+$ cells in uRL. Scale bar indicates 250 μm.

(C) UMAP visualization of GCP sub-clusters from the E16 cerebellum of wild type mice (left panel). Violin plots illustrating the co-expression of *Atoh1* and *Sox2* in slow-cycling progenitors (middle and right panels).

(D) Violin plot showing the potency score in GCP sub-clusters from E16 wild type mice

(E) Violin plots illustrating the number of UMIs and genes in the GCPs of E16 wild type mice.

(F) Heatmap illustrating the differential expression of genes associated with neuronal stem cell maintenance and proliferation, as well as neurogenesis and neuron differentiation, within the GCP sub-clusters from E16 wild type mice.

(G) Bar charts showing the results of the functional enrichment analysis for differential expressed genes in *Sox2*$^+$; *Atoh1*$^+$ cells.

(H) UMAP displaying the presence of *Sox2*$^+$; *Atoh1*$^+$ cells in published cerebellar scRNA-seq datasets.

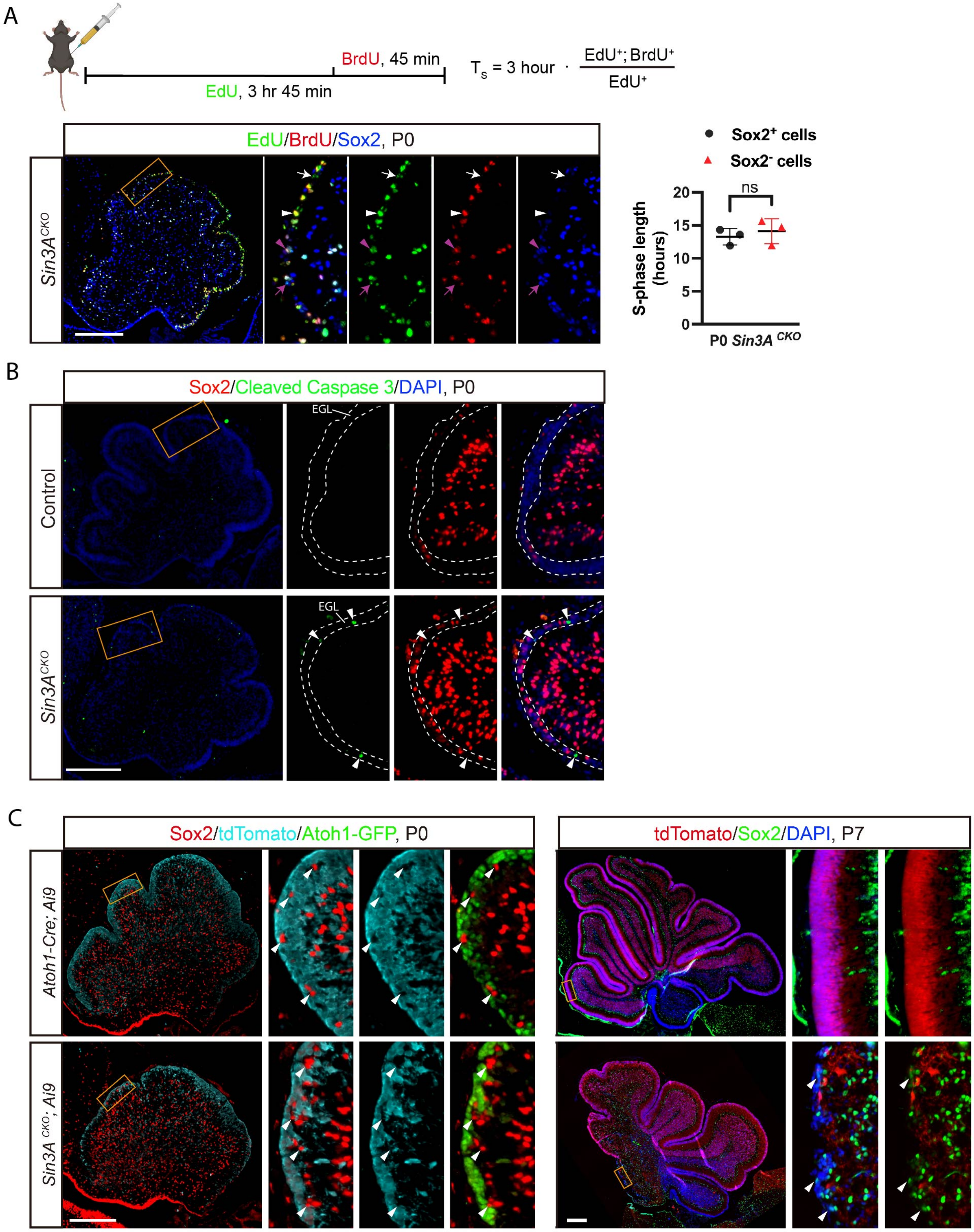
A
BrdU, 45 min
EdU, 3 hr 45 min
$T_s$ = 3 hour · EdU+; BrdU+ / EdU+
EdU/BrdU/Sox2, P0
Sin3A^CKO
Sox2+ cells
Sox2- cells
ns
S-phase length (hours)
20
15
10
5
0
P0 Sin3A CKO
B
Sox2/Cleaved Caspase 3/DAPI, P0
Control
Sin3A^CKO
EGL
C
Sox2/tdTomato/Atoh1-GFP, P0
Atoh1-Cre; Ai9
Sin3A CKO; Ai9
tdTomato/Sox2/DAPI, P7

**Figure S5. Additional images showing characteristics of Sox2$^+$; Atoh1$^+$ cells and Sox2$^+$; Atoh1$^-$ cells in the EGL. Related to Figure 4 and 5.**

(A) The top panel illustrating the strategy for estimating the duration of the S phase using dual labeling with EdU and BrdU. The lower left panel showing Sox2, BrdU and EdU staining in the cerebellum of *Sin3A$^{CKO}$* mouse at P0. White arrowheads indicate Sox2$^-$; BrdU$^+$; EdU$^+$ cells in EGL. White arrows indicate Sox2$^-$; BrdU$^-$; EdU$^+$ cells in EGL. Purple arrowheads indicate Sox2$^+$; BrdU$^+$; EdU$^+$ cells in EGL. Purple arrows indicate Sox2$^+$; BrdU$^-$; EdU$^+$ cells in EGL. Scale bar indicates 250 μm. The lower right panel presenting the quantification of the S phase duration in Sox2$^+$ and Sox2$^-$ cells within the EGL. Shown are mean ± SEM. Two-tailed t test: ns $p > 0.05$. n= 3 mice per group.

(B) Sox2 and CP3 staining in the cerebellum of control and *Sin3A$^{CKO}$* mice at P0. Arrowheads indicate cells positive for CP3. Nuclei were counterstained with DAPI. The dashed lines delineate the boundaries of EGL. Scale bar indicates 250 μm.

(C) Sox2, Atoh1-GFP and tdTomato staining in the cerebellum of control and *Sin3A$^{CKO}$* mice at P0 and P7. Arrowheads indicate Sox2$^+$; tdTomato$^-$ cells in EGL. Nuclei were counterstained with DAPI. Scale bar indicates 250 μm.

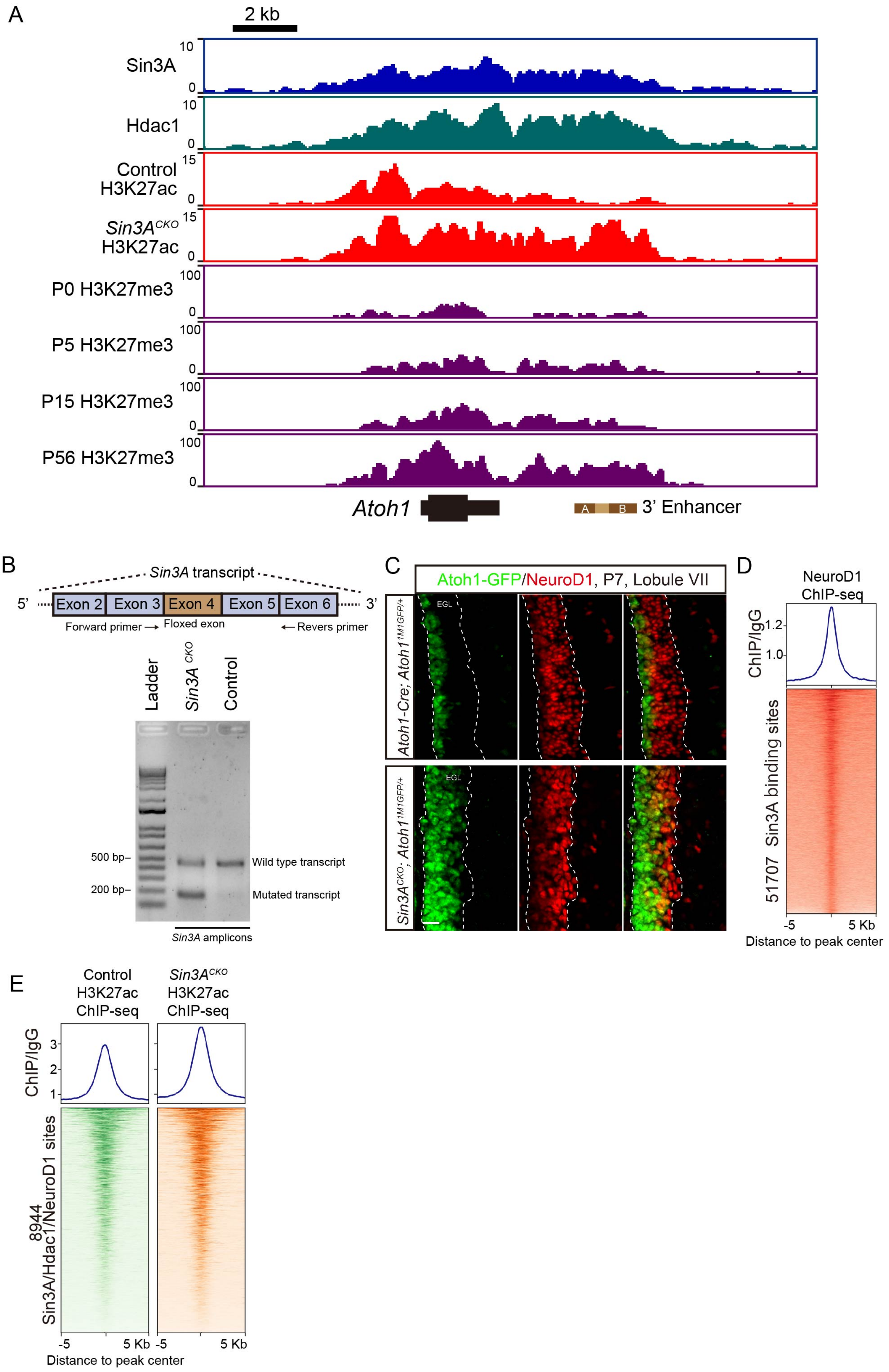

A
2 kb
Sin3A
Hdac1
Control H3K27ac
Sin3A^CKO H3K27ac
P0 H3K27me3
P5 H3K27me3
P15 H3K27me3
P56 H3K27me3
Atoh1
3' Enhancer
B
Sin3A transcript
Exon 2
Exon 3
Exon 4
Exon 5
Exon 6
Forward primer
Floxed exon
Revers primer
Ladder
Sin3A CKO
Control
500 bp–
200 bp–
Wild type transcript
Mutated transcript
Sin3A amplicons
C
Atoh1-GFP/NeuroD1, P7, Lobule VII
Atoh1-Cre; Atoh1 1M1GFP/+
Sin3A CKO; Atoh1 1M1GFP/+
EGL
D
NeuroD1 ChIP-seq
ChIP/IgG
51707 Sin3A binding sites
Distance to peak center
E
Control H3K27ac ChIP-seq
Sin3A^CKO H3K27ac ChIP-seq
ChIP/IgG
8944 Sin3A/Hdac1/NeuroD1 sites
Distance to peak center

**Figure S6. Additional images showing phenotypic changes in the EGL of *Sin3A*$^{CKO}$ mice. Related to Figure 6 and 7.**

(A). ChIP-seq tracks (fold changes of ChIP-seq signal relative to IgG) depicting the intensity of Sin3A, Hdac1, H3K27ac, and H3K27me3 occupation at *Atoh1* locus in the dissociated cerebellar cells or isolated GCPs. The italicized text to the left of each track denotes the genotype of the mice used in ChIP-seq experiments; absence of genotype annotation indicates a wild-type or *Atoh1*$^{M1GFP/+}$ animal. The data for H3K27me$^3$ tracks is derived from previously reported ChIP-seq datasets of the cerebellum at various developmental stages.[110]

(B) RT-PCR results showing the presence of residual wild-type *Sin3A* transcripts in the *Sin3A*$^{CKO}$ GCPs.

(C) Immunofluorescence staining for Atoh1-GFP and NeuroD1 at P6 in the cerebellum of control and *Sin3A*$^{CKO}$ mice. Scale bar indicates 20 μm.

(D) Heatmap and average intensity plots of the NeuroD1 ChIP-seq signals across 10 kb from the center of Sin3A peaks throughout the genome from dissociated cerebellar cells of wild type mice.
